# A disease resistance protein triggers oligomerization of its NLR helper into a hexameric resistosome to mediate innate immunity

**DOI:** 10.1101/2024.06.18.599586

**Authors:** Jogi Madhuprakash, AmirAli Toghani, Mauricio P. Contreras, Andres Posbeyikian, Jake Richardson, Jiorgos Kourelis, Tolga O. Bozkurt, Michael Webster, Sophien Kamoun

## Abstract

NRCs are essential helper NLR (nucleotide-binding domain and leucine-rich repeat) proteins that execute the immune response triggered by disease resistance proteins, also known as sensor NLRs. The structure of the resting state of NbNRC2 was recently revealed to be a homodimer. However, the sensor-activated state has not yet been elucidated. In this study, we used cryo-EM to determine the structure of sensor-activated NbNRC2, which forms a hexameric inflammasome-like structure known as resistosome. To confirm the functional significance of the hexamer, we mutagenized the interfaces involved in oligomerization and found that mutations in three nucleotide-binding domain interface residues abolish oligomerization and immune signalling. Comparative structural analyses between the resting state NbNRC2 homodimer and the sensor-activated homohexamer revealed significant structural rearrangements before and after activation, providing insights into NLR activation mechanisms. Furthermore, structural comparisons between the NbNRC2 hexamer and previously reported CC-NLR pentameric assemblies revealed features in NbNRC2 that allow for the integration of an additional protomer. We also used the NbNRC2 hexamer structure to assess the recently released AlphaFold 3 for the prediction of activated CC-NLR oligomers. This revealed that AlphaFold 3 allows for high-confidence modelling of the N-terminal *α*1-helices of NbNRC2 and other CC-NLRs, a region that has proven difficult to fully resolve using structural approaches. Overall, our work sheds light on the structural and biochemical mechanisms underpinning NLR activation and expands our understanding of NLR structural diversity.

## Introduction

Nucleotide-binding and leucine-rich repeat (NLR) intracellular immune receptors are a key component of innate immunity across the tree of life (1–4). Upon recognition of pathogen derived ligands, they initiate an array of immune responses to counteract infection. In plants, NLRs can be activated by pathogen-secreted virulence proteins, termed effectors, which pathogens deliver into host cells to modulate host physiology (3). A common theme for NLR activation in eukaryotes and prokaryotes is their oligomerization into higher-order immune complexes, such as plant resistosomes or mammalian and bacterial inflammasomes. These complexes initiate immune signalling via diverse mechanisms, often leading to different forms of programmed cell death (1, 5). Some NLRs can function as single units, termed “singletons”, with one NLR protein mediating both ligand/effector perception and subsequent downstream signalling and cell death execution (6). However, NLRs can also function as receptor pairs or in higher order configurations termed NLR networks. In these cases, the paired NLRs have subfunctionalized: one NLR acts as a pathogen sensor that activates another NLR, known as helper, which mediates immune activation and disease resistance (3, 7). While significant progress has been made in recent years regarding the biochemical mechanisms of NLR activation, our understanding of the activation dynamics of paired and networked NLRs remains fragmentary. In particular, our knowledge of the structural biology of helper NLRs is still limited.

NLRs typically exhibit a tripartite domain architecture consisting of an N-terminal signalling domain, a central nucleotide-binding and oligomerization module and C-terminal superstructure-forming repeats (5). In plant NLRs, the central module is termed NB-ARC (nucleotide-binding adaptor shared by APAF-1, plant R proteins and CED-4) and is a defining feature of this protein family (8). The NB-ARC module can be sub-divided into nucleotide binding (NB) domain, helical domain 1 (HD1) and winged-helix domain (WHD) (8). The NB-ARC domain acts as a molecular switch, mediating conformational changes required for transitioning from resting to activated forms (5). In contrast, the N-terminal domain of plant NLRs is variable; it can broadly be used to classify these receptors into distinct groups, which tend to cluster together in NB-ARC-based phylogenetic analyses. In plants, coiled-coil (CC)-type and toll-interleukin receptor (TIR)-type N-terminal domains are the most widespread N-terminal domains (5, 8).

Two plant CC-NLR inflammasome-like structures (resistosomes), AtZAR1 and TmSr35 from Arabidopsis and wheat, respectively (9–11), have been reported to date to be pentameric oligomers. Once activated, these pentameric CC-NLR resistosomes insert into the plasma membrane, mediating Ca^2+^ influx, immune signalling and programmed cell death (10, 12). Importantly, both activated CC-NLR structures correspond to singleton NLRs that do not require other NLRs to function. Our understanding of paired and networked NLRs is comparatively more limited. Activated TIR-type sensor NLRs, such as Roq1 and RPP1, have been reported to assemble into tetrameric resistosomes which activate downstream helper NLRs via small molecule production (13, 14). In contrast, no structures of activated CC-type sensor or helper NLRs have been reported to date. Although activated helper NLRs have recently been reported to oligomerize *in planta*, whether they form tetrameric, pentameric or alternative assemblies is unknown (15–18).

Plant CC-NLRs exhibit diverse N-termini that match the phylogeny of the NB-ARC domain, indicating that they have a deep evolutionary origin (3, 8). To date, two N-terminal CC domains have been defined in angiosperms besides the typical CC domain: RESISTANCE TO POWDERY MILDEW 8 (RPW8)-type (CC_R_) and G10-type CC (CC_G10_) (3, 8). In addition, about 20% or so of the typical CC-type NLRs belong to the wider family of MADA-CC-NLRs, defined by a consensus sequence of their N-terminal *α*1-helices that assemble into the funnel-like membrane pore (19). Both AtZAR1 and TmSr35 can be generally classified as MADA-CC-NLRs. Overall, the N-terminal domain is thought to dictate the types of downstream signalling pathways and activities that take place following NLR activation. However, the structural diversity of these various N-terminal domains and whether they all activate into oligomeric resistosomes in unclear.

Asterids—the largest group of flowering plants—have a complex immune receptor network comprised of a multitude of sensor CC-NLRs that can signal redundantly via downstream helper MADA-type CC-NLRs, the NRCs (NLRs required for cell death) (20, 21). Sensor NLRs in this network include a number of agronomically important disease resistance (R) proteins that function against a variety of microbial pathogens and metazoan pests (21, 22). Previously, our group proposed an activation-and-release model for sensor-helper pairs in the NRC network (16, 17). In this model, effector perception by sensor NLRs leads to conformational changes which expose the central NB domain and enable it to activate downstream NRC helpers (23). This ultimately leads to NRC activation and immune signalling via the assembly of oligomeric resistosomes (15–17, 23). More recently, our group also reported the structure of the resting state of the helper NLR NRC2 from *Nicotiana benthamiana* (NbNRC2), which accumulates as a homodimer prior to activation (24). NRC2 activation by the virus R protein Rx leads to conversion of the helper homodimer into an oligomeric resistosome (24). In parallel, Liu and colleagues reported that a constitutively active mutant of the *N. benthamiana* NRC4 helper, NbNRC4^D478V^, forms a hexameric homo-oligomer with Ca^2+^ channel activity (25). Importantly, whether sensor-activated NRC helpers form hexameric homo-oligomeric assemblies remains to be determined.

In this paper, we report the structure of the sensor-activated helper NLR NbNRC2 purified from the model plant *N. benthamiana* determined by cryogenic electron microscopy (cryo-EM). We show that perception of the *Potato virus X* coat protein (PVX CP) by Rx triggers the formation of homohexameric NbNRC2 resistosomes, distinct from the previously reported pentameric assemblies formed by singleton CC-NLRs. Importantly, the NbNRC2 resistosome does not include the sensor NLR Rx, providing structural evidence for the previously proposed activation-and-release model (17). The structure allowed us to identify and validate the interaction interface between the NbNRC2 protomers in the hexamer, and to pinpoint key residues mediating hexamer assembly. Lastly, we used the NbNRC2 hexamer structure to assess the ability of AlphaFold 3 to predict activated CC-NLR oligomers (26) and showed that it allows for confident modelling of the N-terminal *α*1-helices of NbNRC2 and other CC-NLRs, a feature which has not been well-resolved in most experimental structures determined to date. Overall, our work sheds light on the structural and biochemical mechanisms underpinning NLR activation and expands our understanding of NLR structural diversity.

## Results

### Cryo-EM analysis reveals homohexameric resistosome structure of NbNRC2 activated with Rx and *Potato virus X* coat protein

The *N. benthamiana* helper CC-NLR NbNRC2 exists as a homodimer at resting state and assembles into a higher-order oligomeric resistosome upon activation by Rx or other NRC-dependent sensor NLRs (15–17, 24). The precise structural organization of the activated NRC resistosome is not known. In particular, whether it forms a pentameric assembly that resembles singleton CC-NLRs resistosomes has not been experimentally tested (9–11). To structurally characterize the activated NbNRC2 oligomer, we purified it from *N. benthamiana* following transient overexpression for cryo-EM analysis. We used an NbNRC2 variant with mutations in its N-terminal MADA motif (NbNRC2^EEE^), which was previously identified to abolish cell death induction without compromising receptor activation and oligomerization (17, 19). The C-terminally 3x-FLAG tagged NbNRC2^EEE^ was activated *in planta* by co-expressing the sensor Rx and PVX CP and purified by FLAG-affinity purification (**Fig. S1**). Analysis by negative stain electron microscopy revealed NbNRC2^EEE^ oligomers that were primarily hexameric assemblies. Additional class averages indicated dimerization of the hexamer, producing dodecameric states in a subset of the particles (**Fig. S1A**). Cryo-EM analysis of the sample yielded a reconstruction of the NbNRC2 hexameric resistosome, which was resolved to approximately 2.9 Å resolution with the application of C6 symmetry, into which a structural model was built (**Fig. 1, Fig. S1B-F**).

**Fig. 1:**
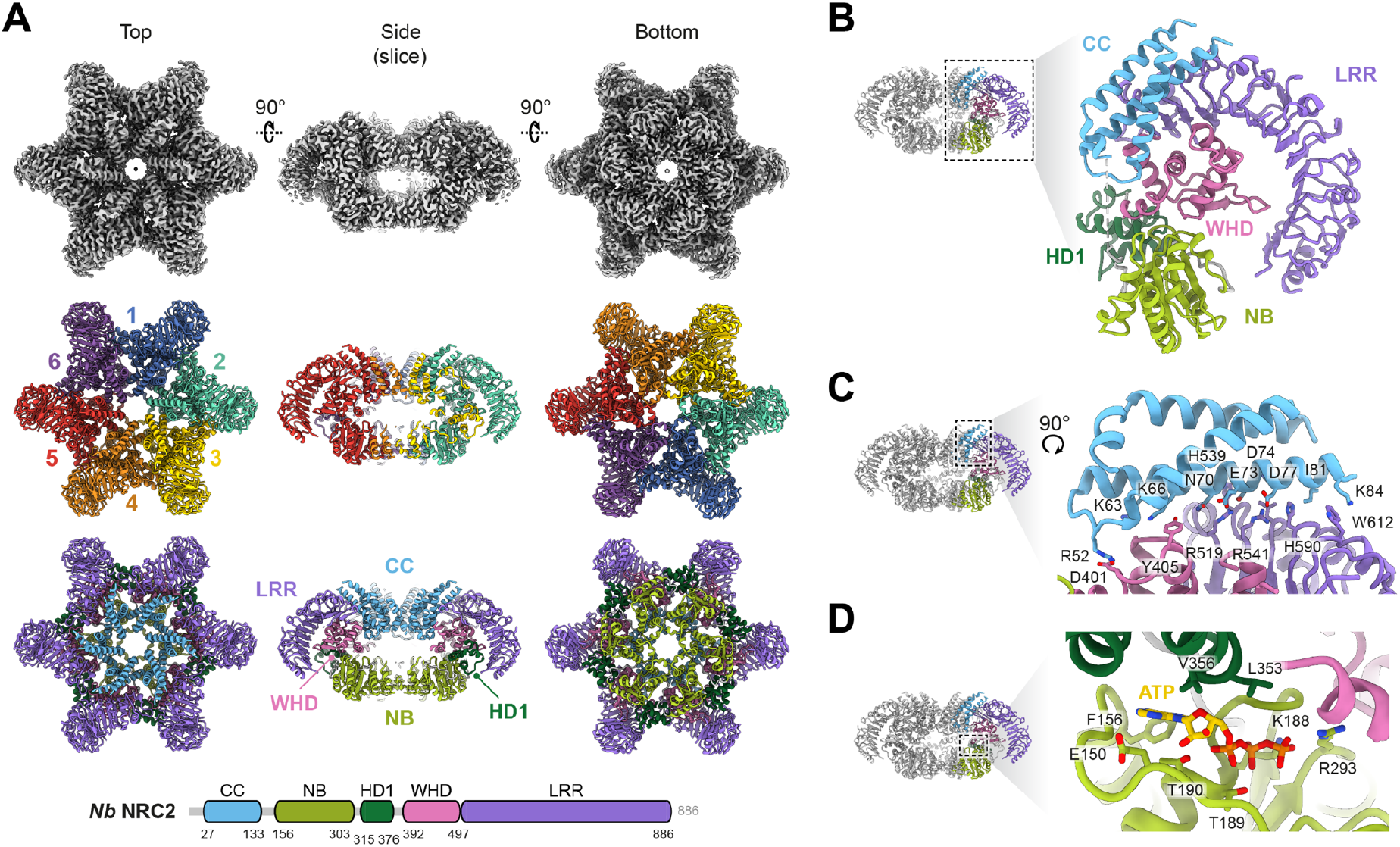
Cryo-EM structure of the NbNRC2 resistosome reveals a homohexameric oligomer. (**A**) The cryo-EM structural analysis revealed that the sensor-activated NbNRC2 assembles into a homohexameric resistosome. The central panel shows the presence of six protomers of NbNRC2. The panel at the bottom displays the individual protomers, coloured by different domains, in accordance with the domain organization (refer to the schematic below) as determined by the structural model. (**B**) For clarity, different domains within a single NbNRC2 protomer and their relative orientations to each other are represented separately. (**C**) The conserved motif EDVID from the α3-helix and its interacting residues from the LRR^R-cluster^ are highlighted, one of the critical contact points important for stabilization of the NbNRC2 resistosome. (**D**) Structural analysis also confirmed the presence of an ATP molecule within the groove formed between the NB domain and HD1 domain of each protomer.

### Overall structure of the NbNRC2 homohexamer

The Rx-activated NbNRC2 oligomer contains six protomers that are assembled into a star-shaped resistosome. The NbNRC2 resistosome therefore contains one more subunit than the previously reported pentameric resistosomes of the MADA-CC-NLRs AtZAR1 and TmSr35 (9–11) (**Fig. 1A**). The *α*1-helices (residues 1-26), which carry the MADA consensus sequence, were not resolved in the NbNRC2 homohexamer, indicating their flexible nature. In contrast, the rest of the CC domain (residues 27-133), NB (residues 156-303), HD1 (residues 315-376), WHD (residues 392-497), and LRR (residues 497-882) regions were resolved (**Table S1, Fig. 1A-B**). Within the NbNRC2 resistosome, the arrangement of domains in each protomer is stabilized by interactions between the CC and LRR domains of the same subunit (**Fig. 1C)**. Specifically, the α3-helix in the NbNRC2 resistosome contacts residues of the LRR and WHD domains. Part of the interaction involves contacts between two conserved motifs that were previously identified: the EDVID motif in the α3-helix and arginine residues from the LRR^R-cluster^ (10, 11). However, the NbNRC2 resistosome also contains features unlike the AtZAR1 and TmSr35 resistosomes. Firstly, NbNRC2 contains an extended EDVID motif in which residues I81, K84, and L85 in the CC α3-helix interact with H590, W612, and Y589, respectively, in the LRR region (**Fig. 1C**). In addition, and unlike the TmSr35 resistosome, only two arginine residues contribute to interactions with the acidic residues from the EDVID motif. Specifically, arginine R519 in the LRR region mediates a bidentate salt bridge with the first two acidic residues of the EDVID motif (E73 and D74) and arginine R541 forms a salt bridge with the final aspartate of the EDVID motif (D77) (**Fig. 1C**). Two potential contacts were observed at the beginning of the α3-helix: residues K66 and N70 likely interact with E495 and N496 in the WHD domain, respectively (**Fig. 1C**).

In the reconstruction, density was observed within the groove formed between the NB domain and HD1 domain of each protomer that is well modelled by a molecule of adenosine triphosphate (ATP) (**Fig. 1D**). The ATP phosphate groups are coordinated by NB domain residues T189, T190, K188 and R293, and the adenine base is within a hydrophobic pocket bordered by F156 and V356 (**Fig. 1D**). The presence of ATP at this position supports its proposed role in stabilization of the resistosome, likely shared with AtZAR1 and TmSr35 (10, 11).

Additional density in the cryo-EM map of the sensor-activated NbNRC2 hexameric resistosome was observed within the concave surface of each LRR repeat around residues W426, K475, K691, N720, K722, K748, and K774 (**Fig. S1G)** which better fits a nucleotide triphosphate. A recent study by Ma et al. on the resting state of SlNRC2 reported inositol phosphate (IP6) or pentakisphosphate (IP5) bound at the same location (27). However, we could not find density corresponding to IP6 or IP5 around these residues (**Fig. S1G**).

### Multiple inter-protomer interactions stabilize the NbNRC2 hexameric resistosome

The NbNRC2 resistosome is stabilized by multiple interfaces between adjacent protomers (**Fig. 2A**). The most extensive interaction that stabilizes the central part of the resistosome is formed by the embedding of each HD1 domain within a concave surface of the adjacent protomer lined by the NB, WHD and LRR regions (**Fig. 2B**). Additional interactions occur between each adjacent CC domain, which form one pore, and each adjacent NB domain, which form the opposing pore. In the contact between CC domains, aspartate D40 in the α2-helix contacts glutamate E60 and lysine K64 of the α3-helix in the adjacent protomer (distances of 3.7 Å and 3.4 Å, respectively) (**Fig. 2C**). In addition, glutamine Q36 of the α2-helix contacts glutamate E60 (distance of 3.8 Å). These extensive electrostatic interactions between CC domains likely stabilize their position within the NbNRC2 resistosome.

**Figure 2:**
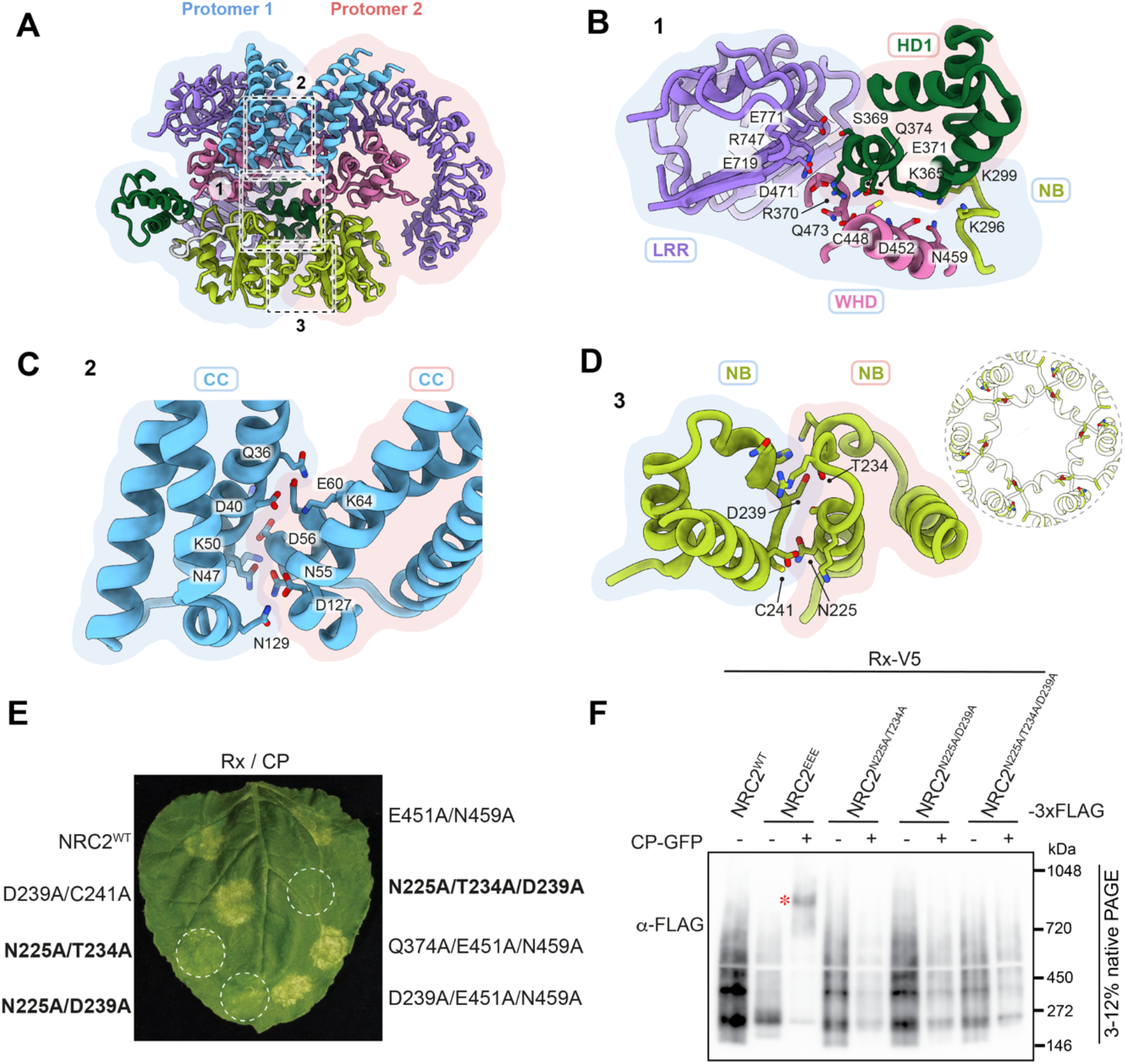
I*nter*-protomer interactions stabilize the NbNRC2 resistosome. **(A)** Multiple potential contacts between the two protomers of the NbNRC2 resistosome, highlighted in dashed boxes, typically contribute to its overall stability. (**B**) The HD1 domain from one protomer establishes extensive interactions with the NB, WHD and LRR domains of the adjacent protomer, stabilizing the NbNRC2 resistosome. (**C**) CC *vs.* CC interactions between the two adjacent protomers specifically stabilize the NbNRC2 homohexamer from the topside. (**D**) Similarly, residues at the NB *vs.* NB interface contribute to the stabilization of the resistosome from the underside. The inset shows the NB pore of the NRC2 resistosome and how it is well connected by the four crucial amino acids. Potential contact residues from all these interfaces are highlighted in stick representation. (**E**) A representative leaf picture from *N. benthamiana nrc2/3/4* KO plants showing HR after co-expression of Rx/PVX CP with NRC2^WT^ and its combinatorial variants. (**F**) BN-PAGE assay with inactive and activated Rx with NRC2^EEE^ and the NRC2^WT^ mutants, which exhibited weak or no hypersensitive cell death. Sensor-helper combinations shown were co-infiltrated together with GFP or CP-GFP. Total protein extracts were run in parallel on native and denaturing PAGE and immunoblotted with the antisera labelled on the left. SDS-PAGE blots are found in **Fig. S4**. Approximate molecular weights (in kDa) of the proteins are shown on the left. An asterisk represents the higher-order oligomer state for NRC2^EEE^ upon activation with Rx and PVX CP. The experiment was repeated three times with similar results.

At the interface between NB domains, aspartate D239 and cysteine C241 from one protomer contact threonine T234 and asparagine N225, at distances of 3.3 Å and 4.4 Å, respectively, in the other protomer (**Fig. 2D**). To assess the contribution of the contact between NB domains we introduced mutations into key residues at the interface and assayed the mutants for activation of immunity and resistosome formation. NbNRC2 variants with single mutations in the inter-protomer interface were unaltered in their ability to cause hypersensitive cell death (**Fig. S2**). From a set of seven combinatorial mutations, we identified two variants that contained mutations of two residues that showed weak cell death activity following activation by Rx and PVX CP (NbNRC2^N225A/T234A^ and NbNRC2^N225A/D239A^), and a variant with mutation of three residues that displayed an almost complete loss of cell death activity (NbNRC2^N225A/T234A/D239A^) (**Fig. 2E, Fig. S3**). The protein abundance of double and triple NbNRC2 mutants were comparable to NbNRC2^WT^, indicating that the lack of cell death is not due to these NbNRC2 variants accumulating poorly (**Fig. S3**). As the cell death response was compromised, we were able to assess the oligomerization of the mutants upon Rx-mediated activation using BN-PAGE assays, as previously reported (16, 17). Unlike NRC2^EEE^, the double mutants and the triple mutant did not form detectable higher-order resistosome-like complexes upon Rx-mediated activation (**Fig. 2F**).

From these results, we conclude that the residues at the NB domain interface, N225, T234, and D239 are critical to resistosome formation. It is notable that mutation of a small number of residues within the NB domain is sufficient to compromise hexamer formation given the extensive inter-protomer interface that involves all domains of NRC2 (**Fig. 2B**). We conclude that contact between NB domains plays a significant role in the oligomerization pathway and that the more extensive interactions between other domains are not stable in its absence. Furthermore, the contribution of the interaction between NB domains is comparable to the role of the interaction between the well-conserved EDVID motif from the α3-helix and the LRR^R-cluster^ at the topside of the resistosomes.

### Comparison between resting state homodimer and activated hexamer of NbNRC2 reveals extensive NB-ARC conformational rearrangements

Our model allows us to examine the mechanism of resistosome formation by comparing the active hexameric resistosome to that of the dimeric resting state. We performed a comparative structural analysis assessing the conformation of NbNRC2 protomers in each state and the interactions between them that support oligomerization. We identified that NbNRC2 contains two structural modules that vary in their relative position between the hexameric and dimeric states (**Fig. 3A**). The first structural module comprises the NB and HD1 domains and the second comprises the WHD and LRR domains. The relative orientations of the domains within each module are mostly unchanged between the hexameric and dimeric states. By contrast, the two modules must undergo a rotation of approximately 180° with respect to each other to transition between the states (**Fig. 3B, Movie S1**).

**Fig. 3:**
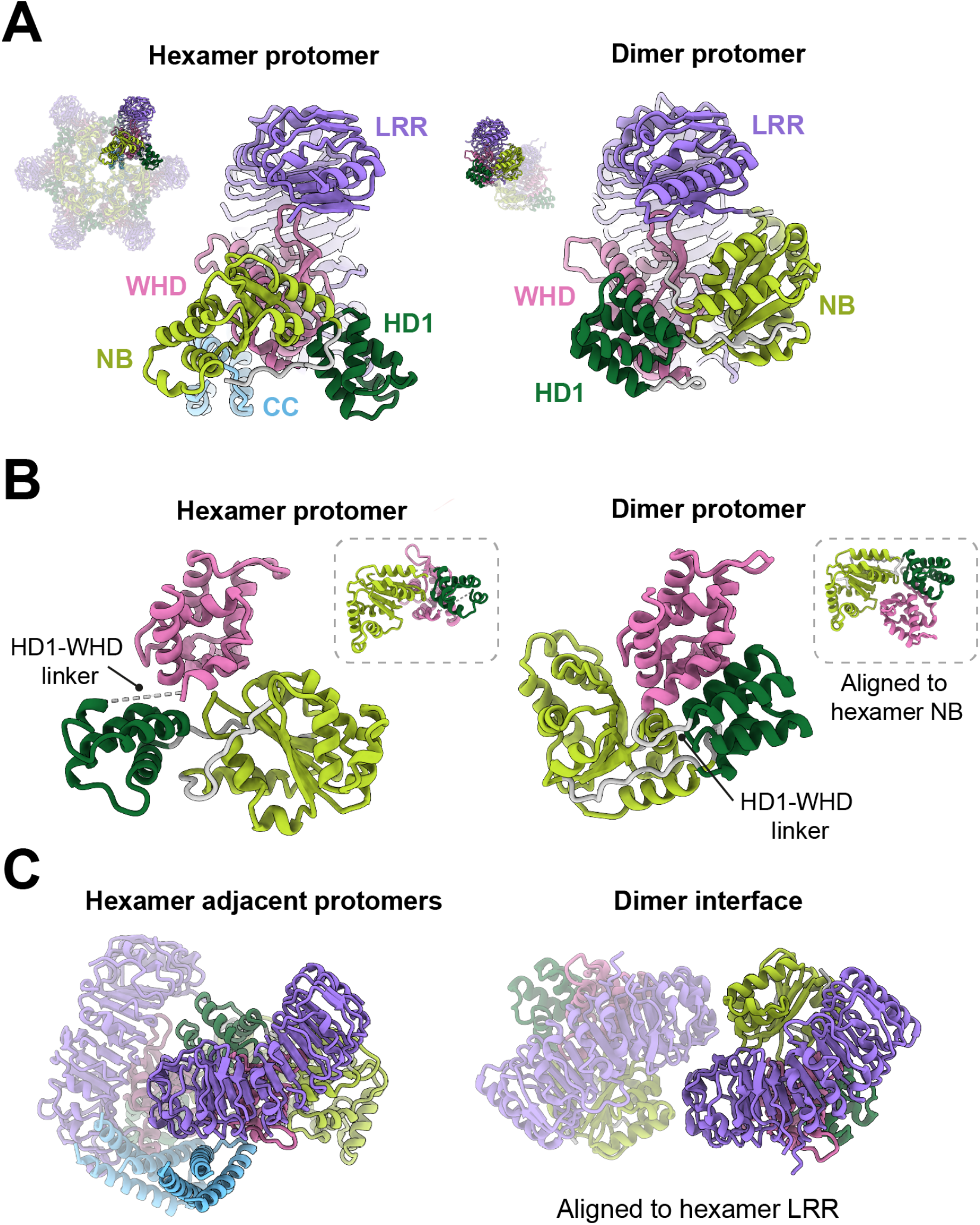
A protomer-level structural comparison between the NbNRC2 hexamer and dimer reveals major differences in the NB-ARC regions. (**A**) A comparison of the individual protomers from the NbNRC2 hexamer (PDB ID: 9FP6) and dimer (PDB ID: 8RFH) complexes, revealed crucial differences in the relative orientation of the NB, HD1 and WHD domains within the NB-ARC regions. The difference in the relative orientation of the NB and WHD domains, along with the contribution from the HD1-WHD linker region to this switch, was further highlighted by aligning to the hexamer NB (**B**) and hexamer LRR region (**C**), independently. The missing density for the HD1-WHD linker from the hexamer protomer is indicated by a dashed line.

The transition between resting dimeric and active hexameric states involves additional changes. Firstly, the linker between the HD1 and WHD domains that connects the structural modules is ordered in the dimeric state but disordered in the hexamer. Secondly, the extensive interactions between the WHD, HD1 and NB domains of a single protomer in the dimeric state must be disrupted to allow the NB-HD1 module to rotate to a new position with respect to the WHD-LRR module (**Fig. 3B, Movie S1**). Thirdly, the CC domain is resolved in the hexamer but not the dimer state suggesting that it is disordered or highly flexible in the dimeric state but partly ordered in the hexamer, except for the *α*1-helix. Finally, the surfaces that form inter-protomer interaction interfaces in the NbNRC2 homodimer are not in contact in the hexamer. The dimer interface involves contacts between the LRR and NB domains and these domains have different orientations in the hexamer (**Fig. 3C**) (16). We conclude that the characterized homodimeric state undergoes significant conformational rearrangements in order to form the active hexamer upon Rx-mediated activation, suggesting that intermediate conformations may occur.

### The NbNRC2 hexamer exhibits structural commonalities and differences with the previously reported MADA-CC-NLR pentameric resistosomes

NbNRC2 is a member of the MADA class of CC-NLRs like AtZAR1 and TmSr35. The discovery that NbNRC2 forms a hexameric state upon activation, as recently reported for NRC4^D478V^ (25), reveals unexpected diversity in MADA-CC-NLR oligomerization given that AtZAR1 and TmSr35 are both pentameric. To investigate the structural basis of hexamer rather than pentamer formation we compared the structures of characterized CC-NLR resistosomes (9–11). Despite significant difference in overall resistosome architecture caused by the accommodation of an additional subunit, the conformation of individual NbNRC2 protomers closely resembles those of AtZAR1 and TmSr35 (**Fig. 4A**). In each resistosome, corresponding regions of the CC and NB domains support inter-subunit interactions but the identity of residues at the interfaces vary considerably.

**Fig. 4:**
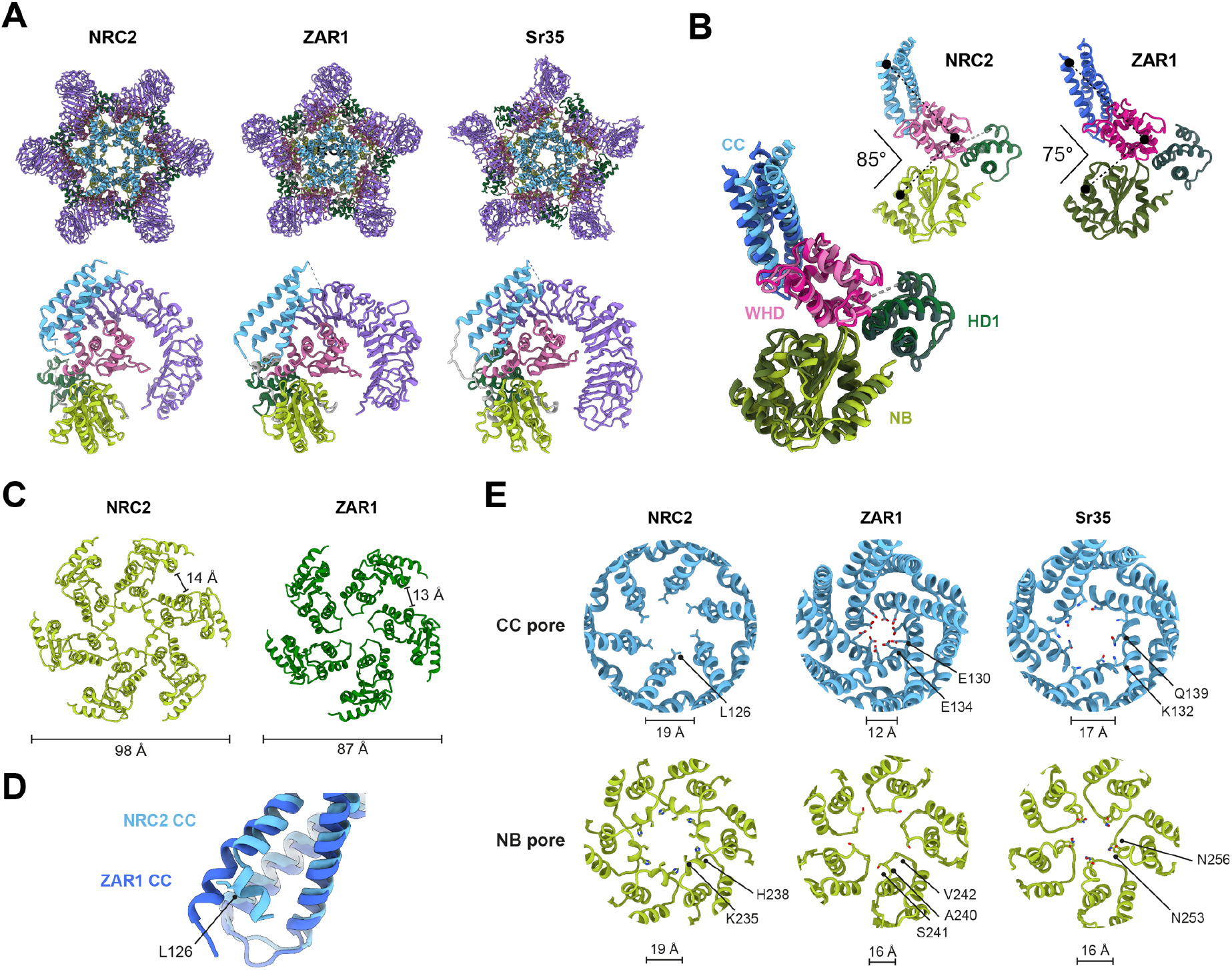
Comparison of NbNRC2, AtZAR1, and TmSr35 resistosomes reveals striking structural differences. **(A)** The sensor-activated NbNRC2 assembles into a homohexameric resistosome, in contrast to AtZAR1 and TmSr35, which form pentameric resistosomes. The induvial protomers (coloured by domains) from these resistosomes still share the same topology. (**B**) Structural superposition of the protomers from the NbNRC2 hexamer and the AtZAR1 pentamer revealed differences at the protomer level. Specifically, the angular distances between the domains within the individual protomers differ, potentially influencing the packing of protomers within the respective resistosomes. (**C**) Further, this arrangement results in an overall outward displacement of the ‘NB’ domains, creating a wider underside in the NbNRC2 resistosome, while maintaining a nearly constant distance between the domains. (**D**) Structural superposition of the NbNRC2 and AtZAR1 CC domains revealed a ‘kink’ in the α4-helix of the CC domain in the individual protomers from NbNRC2 resistosome. (**E**) A comparison of the CC and NB pores, and the chemical nature of the pores, in the NbNRC2, AtZAR1, and TmSr35 resistosomes. Distances indicated are measured between the C^α^ atoms of the two opposite residues highlighted in the respective resistosomes.

The relative positions of the domains that comprise the NB-ARC module differs between NbNRC2 and AtZAR1 in a way that is consequential to the stoichiometry of the complex (**Fig. 4B**). The interdomain angle within the CC-NB-ARC was measured to be 10° larger in NbNRC2 relative to AtZAR1. This widening of the NB-ARC has two consequences. Firstly, the NB domains are pushed outward in NbNRC2 relative to their positions in AtZAR1 (**Fig. 4C**). This difference is significant for the accommodation of an additional subunit in the NbNRC2 resistosome. Secondly, the cavity bordered by the LRR, WHD and NB on the internal face of the protomer that interacts with the HD1 domain of the adjacent protomer (**Fig. 2B**) is substantially larger in NbNRC2 than AtZAR1. This allows the HD1 domain to insert further into the cavity and make more extensive contact. Finally, a structural difference in the CC domain also contributes to the stoichiometry of the resistosome. The α4-helix of NbNRC2 contains a bend near residue L126, which is at the end closest to the resistosome core (**Fig. 4D**). This difference is significant as structural superposition showed the straight α4-helix of AtZAR1 would sterically overlap with the adjacent protomer in a hexameric configuration.

The presence of an additional protomer and the increased separation of the NB domain positions prompted us to examine whether the pores of the NbNRC2 resistosome are larger than those of pentameric resistosomes. We identified that NbNRC2 and TmSr35 have a one third wider CC pore (17–19 Å) compared to the AtZAR1 (12 Å) (**Fig. 4E**). The small CC pore of AtZAR1 is a consequence of the α4-helix, which is longer than that of TmSr35 and unbent unlike NbNRC2. On the opposing side, the NB pore of all three resistosomes has a similar width, though that of NbNRC2 is 3 Å larger (19 Å) than those of AtZAR1 and TmSr35 (16 Å) (**Fig. 4E**). Thus, the stoichiometry of the resistosome is not the only significant factor in determining pore diameter. We additionally observed that the residues that line the pores of each resistosome are different across characterized resistosomes. Whereas the CC pore of NbNRC2 is hydrophobic due to a ring of leucine residues (L126), the CC pore of AtZAR1 is acidic due to a ring of glutamate residues (E130 and E134) and the CC pore of TmSr35 is of mixed charge (K132 and Q139). We conclude that the size and chemical composition of resistosome pores is unexpectedly variable, with potential consequences for their channel activity.

### AlphaFold 3 can predict a lipid-bound NbNRC2 resistosome with high confidence

Activated CC-NLR resistosomes insert themselves into membranes via a funnel-like structure formed by their N-terminal α1 helices, leading to the induction of cell death. The absence of the α1-helix and the incomplete pore in the NbNRC2 resistosome cryo-EM structure led us to use AlphaFold to generate a structural prediction for the missing region to obtain an integrated model that is more complete (26). First, we used the experimentally resolved NbNRC2 hexamer structure to assess the performance of AlphaFold 2, AlphaFold 3 and AlphaFold 3 with 50 oleic acid molecules as a stand-in for the plasma membrane (26, 28, 29). We focused on modelling the CC-NB-ARC portion of the full-length NbNRC2, which includes the oligomerizing domains. We analyzed the models based on different metrics such as predicted local distance difference test (pLDDT), predicted template modelling score (pTM), interface predicted template modelling (ipTM), predicted aligned error (PAE), minimum per-chain pTM, and chain-pair ipTM (26, 29). AlphaFold 2, AlphaFold 3 and AlphaFold 3 with lipids, all generated high-confidence models with pTM and ipTMs of 0.7 and higher and structurally aligned with the experimental model with RMSD values of lower than 1.8 Å (**Fig. 5A, Fig. S5A, Fig. S5B**). However, in the AlphaFold 2 model, the N-terminal α1-helix faced inwards, whereas in both AlphaFold 3 predicted structures the N-terminal α1-helix was facing outwards and formed a full funnel-shaped structure (**Fig. S5A, Fig. S5A, Fig. S5B**). In the AlphaFold 3 model with 50 oleic acids, the N-terminal α1-helix showed confident interactions with the lipids and had a higher local confidence compared to the models without the lipids (**Fig. 5B, Fig. S5A, Fig. S5B**).

**Fig. 5.**
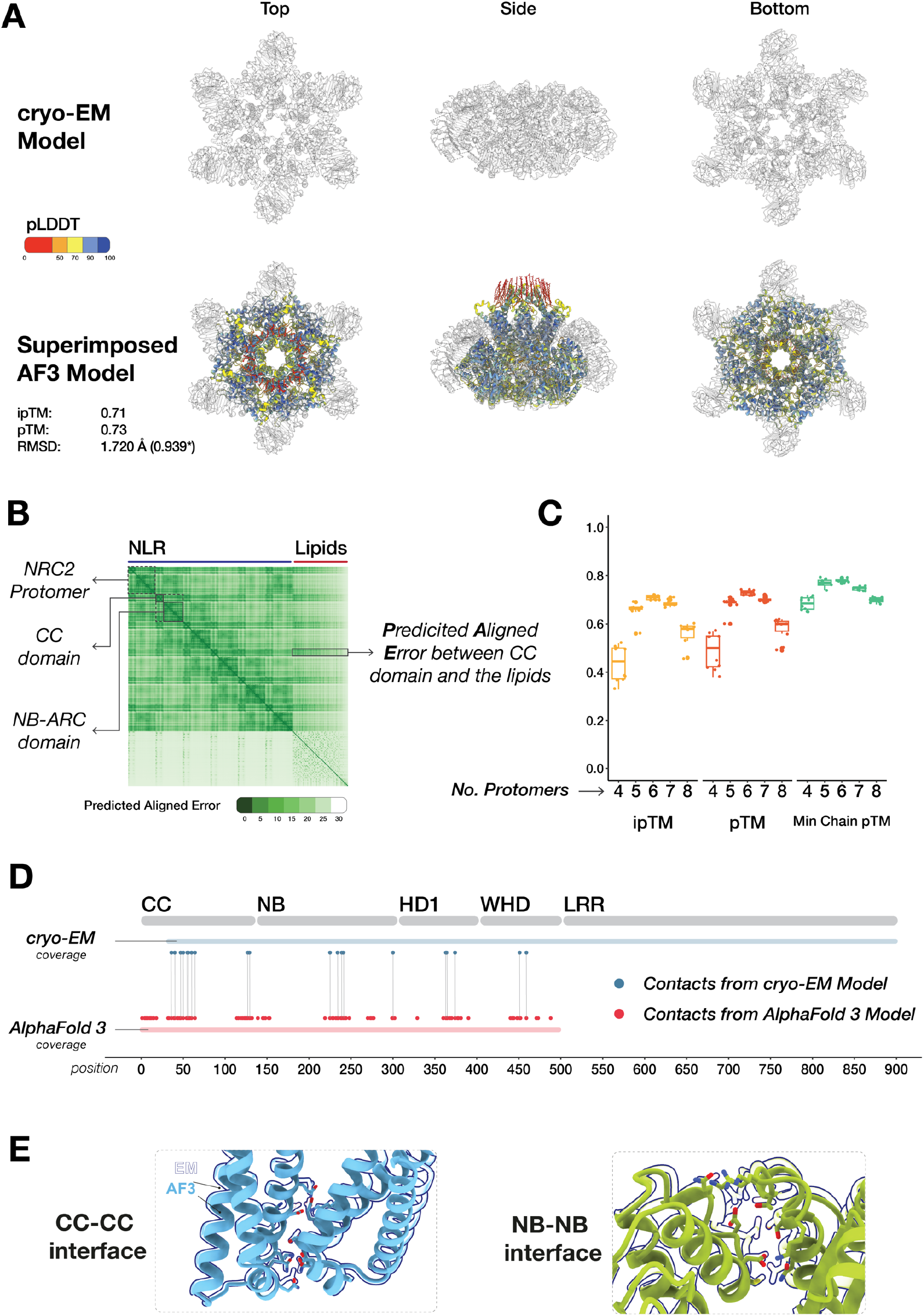
AlphaFold 3 can predict a high-confidence NbNRC2 resistosome. (A) Superimposition of NbNRC2 predicted model with the cryo-EM structure. NbNRC2 AlphaFold 3 resistosome model fits into the cryo-EM structure (PDB ID: 9FP6) with an RMSD of 1.72 Å. Structures are colored by pLDDT values. (**B**) Predicted aligned error plot of NbNRC2 AlphaFold 3 model. Representative single protomers, domains, and interactions between the CC-domain and the lipids are shown. (**C**) ipTM, pTM, and Minimum Chain pTM confidence values of NbNRC2 predicted oligomers (N = 10). (**D**) NbNRC2 AlphaFold 3 resistosome model comprises overlapping contact regions as the cryo-EM structure. (**E**) Predicted NbNRC2 model superimposed on selected contact regions in the cryo-EM model.

How does varying the NbNRC2 stoichiometry affect the confidence of the AlphaFold 3 predictions? To assess this, we repeated the process using a range of stoichiometries from 4 to 8 NbNRC2 copies. For each stoichiometry, we generated 10 models with different seeds each time, all modelled with 50 oleic acid molecules (**Fig. 5C**). Hexamer models had the highest median pTM and ipTM confidence scores and aligned with the experimental NbNRC2 structure with RMSD values lower than 1.8 Å (**Fig. 5C, Fig. S6**). Predictions for pentamers and heptamers were less confident across all metrics, while tetramers and octamers were of lower confidence and showed greater variability (**Fig. 5C**). All oligomeric forms consistently displayed complete funnel-like structures with the exception of tetramers. Tetramer models varied, with some forming complete funnels and others resembling incomplete hexameric-like resistosomes (**Fig. S7A**).

To further assess the AlphaFold 3 models, we checked if the contact residues from the cryo-EM structure were also predicted in the AlphaFold 3 resistosome structures. Based on a 4.5Å cut-off for contact, most residues involved in interactions in the cryo-EM model were also predicted in the AlphaFold 3 model (**Fig. 5D, Data S2**). However, the ability to predict the details of the interaction interface varied. While the interaction between adjacent CC domains was predicted approximately correctly, the residues at the interface between adjacent NB domains were dissimilar to those in the experimental structural model (**Fig. 5E, Fig. S7B, Fig. S7C**).

### AlphaFold 3 prediction of CC-NLR resistosomes reveals a diversity of N-terminal pore-like structures

The ability of AlphaFold 3 to predict the NbNRC2 resistosome stoichiometry and architecture with confidence prompted us to examine predicted models of other NLR proteins. We first modelled 11 representatives from the major NRC protein clades as pentamers and hexamers (29). 3 of the 11 NRC proteins didn’t show confident models with SlNRC0, NbNRCX, and NbNRC7a having ipTM and pTM scores lower than 0.6 as both pentamers and hexamers (**Fig. S8, Fig. S9**). The remaining 8 NRCs all had confidence values of 0.6 and higher for both pentamer and hexamers (**Fig. S8, Fig. S9**). In all 11 cases, hexamers had higher confidence values in all models compared to pentamers, although this difference was subtle (**Fig. S9**). Additionally, all hexamer models except NbNRC7a had minimum per-chain pTM scores of 0.7 and higher. In contrast, only eight out of twelve NRCs had minimum per-chain pTM scores of 0.7 and higher as pentamers (**Fig. S9**).

We also determined whether AlphaFold 3 can model the CC-NB-ARC portion of AtZAR1 and TmSr35, the two CC-NLRs with experimentally determined pentamer resistosome structures (9–11). We used stoichiometries of either 5 or 6 copies. Although not predicted with as high confidence as NbNRC2, the predicted pentamer models of AtZAR1 and TmSr35 aligned well to experimental models, with RMSD values of 1.511 Å and 1.76 Å respectively (**Fig. 6A**). Each NLR had higher confidence for the pentameric form compared to the hexameric form, reflecting their experimentally validated resistosome configurations (**Fig. S10**).

**Fig. 6.**
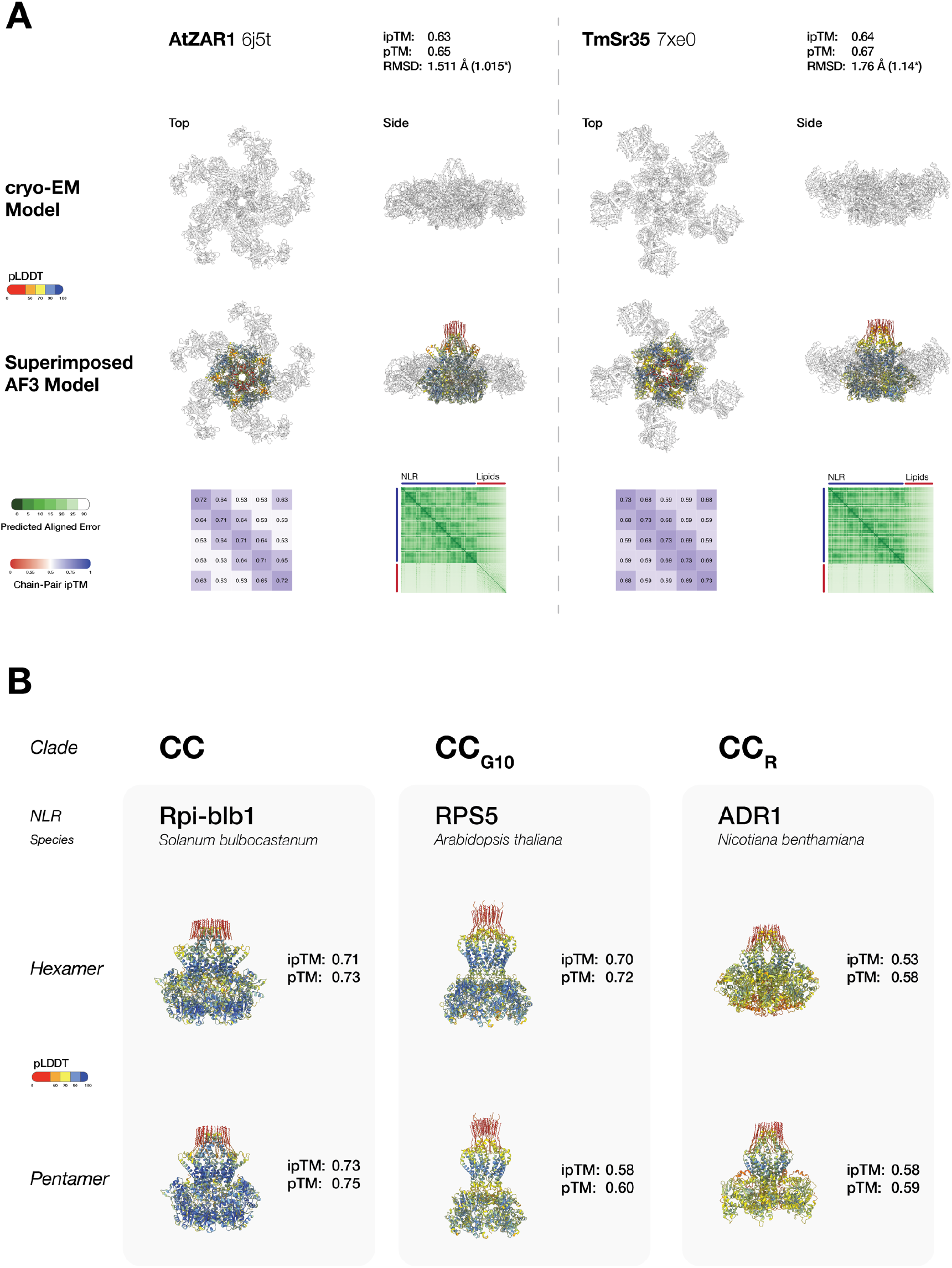
Other CC-type NLR resistosomes can be predicted confidently using AlphaFold 3. (**A**) Superimposition of AtZAR1 and TmSr35 AlphaFold 3 predicted resistosome models on their available cryo-EM structures (PDB IDs: 6J5T, 7XE0) with predicted align error and chain-pair ipTM confidence plots. (**B**) Predicted AlphaFold 3 resistosome structures of SbRpi-blb1, AtRPS5, and NbADR1 as pentamers and hexamers. Structures are colored by pLDDT values.

Given AlphaFold 3 performance in modelling CC-NLRs such as NbNRC2, AtZAR1, and TmSr35, we proceeded to model a set of phylogenetically diverse plant NLR proteins with unknown resistosome structures (**Fig. S11, Data S1**). We chose 8 CC-type, 6 CC_G10_-type, and 2 CC_R_-type NLRs and modelled them with AlphaFold 3 as pentamers and hexamers to assess if their resistosome structures could be predicted with high confidence metrics and to determine which oligomeric configuration yielded better models (**Fig. 6B**).

This approach revealed that the CC-type and CC_G10_-type NLR clades each contain representatives with hexamer and pentamer resistosome configurations. In the CC-NLR clade, all NLRs modelled with ipTM, pTM, and minimum chain pTM values of 0.6 or higher (**Fig. S12-Fig. S14, Fig. S18**). AtHrt1 was confidently predicted to be a hexamer, whereas AtRpm1 and SlPtr1 were more confident as hexamers than pentamers. OsPit, TaPm2a, StR3a, SbRpi-blb1, and BvRz2 all showed higher model confidence for pentamers (**Fig. S12-Fig. S14**). However, in StR3a and BvRz2, the CC domain and the N-terminal α-helix were modelled with lower confidence than in the other resistosome models in either oligomeric form (**Fig. S13, Fig. S14**).

CC_G10_-NLRs showed variable model confidence for hexamers and pentamers. AtRPS5 had higher confidence as a hexamer (**Fig. S16**), whereas CaPvr4, LsRGC2B, GmRsv3, and CmVat showed higher model confidence in their pentameric form (**Fig. S15, Fig. S16**). Compared to the rest of the CC_G10_-NLRs, AtRPS2 was poorly modelled in both stoichiometries, with confidence values below 0.6 (**Fig. S15, Fig. S18**).

The two CC_R_-NLRs modelled, NbADR1 and NbNRG1, did not yield high-confidence models as pentamers or hexamers, with confidence values below 0.6 in all cases (**Fig. S17, Fig. S18**). However, while the overall confidence values were low, they showed higher confidence as pentamers compared to hexamers.

## Discussion

Here, we report the structure of a sensor-activated plant helper NLR determined by cryo-EM. We report that, following perception of the coat protein of *Potato virus X* by the disease resistance protein Rx, the helper CC-NLR NbNRC2 assembles into a hexameric resistosome that is different to the pentameric complexes formed by activated singleton MADA-type CC-NLRs (**Fig. 1**). The NbNRC2 resistosome does not contain Rx, providing structural evidence for the activation-and-release mechanism for sensor-helper pairs in the NRC immune receptor network (17). This indicates that this mode of activation of paired NLRs is markedly distinct from the hetero-complexes of metazoan paired NLRs, such as NAIP/NLRC4 (17, 30). We conclude that paired and networked inflammasomes and resistosomes from various biological systems assemble through diverse mechanisms (2, 30).

Our study provides insights into helper NLR activation by sensors, which involves oligomerization into a hexameric resistosome. This expands our current understanding of NLR structural diversity beyond tetrameric and pentameric activated oligomers. What is the functional relevance of the NbNRC2 hexamer? In previous studies, we showed that activated NbNRC2 and NbNRC4 oligomers accumulate as plasma membrane-associated puncta following activation by sensors, leading to cell death and disease resistance (17, 31). Recently, the NbNRC4^D478V^ oligomer was also shown to function as a calcium channel as has been shown for pentameric resistosomes of CC-NLRs AtZAR1 and TmSr35 (10, 12, 25). Considering that the NbNRC2 hexamer features a CC pore size that is one third larger than AtZAR1, as well as different inner volumes (**Fig. 3**, **Fig. 4**), it will be interesting to understand if these differences lead to altered ion channel dynamics compared to CC-NLR pentamers. Moreover the extent to which the NbNRC2 hexamer serves as a channel for influx/efflux of other unknown immunogenic small molecules that cannot be channelled by smaller, pentameric assemblies remains to be determined. The relevance and consequences of resistosome pore size on downstream immune activation will be an exciting question to answer.

Interestingly, while we identified density corresponding to a nucleotide triphosphate around residues W426, K475, K691, N720, K722, K748, and K774 in the active NbNRC2 resistosome, a recent study by Ma et al. reported inositol phosphate (IP6) or pentakisphosphate (IP5) bound at the same location in the resting state structure of SlNRC2 (27). Mutation of these residues in SlNRC2 resulted in either reduced or complete loss of cell death or conductivity upon activation with Rx and PVX CP (27). Whether or not mutations at these positions affect hexamerization remains to be tested.

The NbNRC2 hexamer is the fourth NLR to be structurally characterized by cryo-EM using purification from the model plant *N. benthamiana*, underscoring the versatility of this emerging expression system for structural biology. The capacity to purify NLR protein complexes directly from plants and characterize them using cryo-EM promises to accelerate our understanding of immune receptor structural diversity (13, 24, 25). Importantly, whether other CC-NLRs assemble into hexamers remains to be determined. Indeed, it is tempting to speculate that additional resistosome stoichiometries remain to be discovered.

Of note, the pentameric CC-NLR structures solved to date were obtained from samples purified from insect cells and not plants (10, 11). It therefore remains to be shown whether the expression system can have an impact on the order of oligomer formed. However, AlphaFold 3 predictions of AtZAR1 and TmSr35 with different stoichiometries yielded higher confidence for pentameric assemblies (**Fig. 6A**), suggesting that the pentamers are apparently not artifacts of insect cell expression. In the future, it would be important to purify these complexes directly from plant tissue and compare them to the protein complexes produced in insect cells.

Comparative analyses of the resting and activated forms of NbNRC2 revealed distinct interfaces involved in homodimerization compared to homohexamerization and extensive intramolecular rearrangements within each NbNRC2 protomer, particularly in the NB-ARC domain (**Fig. 3**). It will be interesting to determine whether there are subtle differences between the overall structure and the conformational changes of individual domains between NbNRC2 and other NRC hexamers (25). In the NbNRC2 hexamer, the homodimerization interface is largely surface exposed, clearly suggesting that this interface needs to be disrupted in order to allow for resistosome formation (**Fig. 3, Movie S1**). However, whether its disruption leads to homodimer dissociation into individual monomers which then hexamerize is not clear. Importantly, we could not detect the appearance of a monomeric NRC2 species in our BN-PAGE assays using hexamerization interface mutants unable to oligomerize (**Fig. 2**). Another alternative is that the NbNRC2 homodimer remains associated upon activation, but that the individual protomers in the homodimer undergo extensive conformational rearrangements that allow them to oligomerize. Indeed, three primed homodimers could assemble into a hexamer, and this could explain the absence of pentameric NRC2. A mechanism that involves trimerization of homodimers would provide a straightforward explanation for hexamer assembly. Future experiments will shed light on the precise dynamics of NRC homodimer conversion into hexamers.

So far, plant NLRs have been shown to assemble into tetrameric, pentameric and now hexameric resistosomes (3, 5). That the highly conserved NB-ARC domain of plant NLRs can form oligomeric assemblies of such different stoichiometry is intriguing and highlights its flexible nature. Further analyses, possibly assisted by AlphaFold 3, may help identify the molecular features within the NB-ARC and CC domains that determine resistosome stoichiometry, which may potentially allow for predicting the number of protomers in an activated NLR resistosome based solely on amino acid sequence. Similarly, we previously showed that the AtZAR1 *α*1-helix can functionally complement cell death activity and disease resistance when swapped into NRCs (19, 32). That the *α*1 helix from a pentamer-forming CC-NLR can still function just as well in the context of a hexamer-forming oligomer indicates that these N-terminal helices encode features at the amino acid level which allow them to be quite versatile and capable of forming funnels with different number of units and pore sizes.

We found that AlphaFold 3 can predict the overall architecture of the NbNRC2 hexamer and a subset of the interaction interface residues with high confidence (**Fig. 2**, **Fig. 5**). Moreover, we found that the inclusion of oleic acids in the prediction allowed it to confidently predict the N-terminal funnel formed by the *α*1-helices, which is not always resolved in experimental CC-NLR resistosome structures (**Fig. 5, Fig. S7C**) (26). We hypothesized that this approach could be effective for predicting membrane-bound protein structures as recently proposed (28). Our assessment, using three experimental CC-NLR resistosome structures revealed that AlphaFold 3 can predict activated NRC hexamers and CC-NLR pentamers with high confidence. More importantly, including oleic acids in the prediction allowed for modelling of the N-terminal funnel formed by the *α*1-helices of the CC-NLR protomers with high confidence. Using this approach, we obtained confident predictions of N-terminal helices for a representative set of NLRs with coiled coil N-termini, including canonical CC type and CC_G10_-type NLRs. This suggests that these NLRs are likely to assemble into resistosomes with diverse pore-like N-terminal structures. However, it’s worth noting that CC_R_-NLRs and several CC-NLRs modelled did not yield high confidence resistosome models. The full significance of these predictions will require further investigations.

The N-terminal funnels of activated CC-NLR resistosomes are often not resolved in experimental structures, either due to the mutations introduced in the *α*1-helices to abolish cell death or due to depletion of lipids during purification, which may play a key role in stabilizing these N-terminal funnels (9, 10, 25). In this sense, AlphaFold 3 can be used to fill out gaps in resistosome structures that have remained elusive due to technical limitations. More importantly, these results point to a new potential application for AlphaFold 3 as a tool to study the structural diversity of activated NLRs, serving as a platform for hypothesis generation and NLR classification without the need for experimental structures. Indeed, in ongoing work in our group, we have been using AlphaFold predictions to complement phylogenetic grouping and other data to triage NLRs into functional categories and prioritize them for functional analyses.

Ever since the first plant NLR resistosome structure was solved in 2019 (11), a multitude of key structural, biochemical and cell biology studies have resulted in models for NLR activation, largely based on the initial AtZAR1 resistosome (3, 5). This present study together with our recent work on the NbNRC2 resting state homodimer (24), expands the current NLR paradigms beyond the monomer to pentamer model established for singleton CC-NLRs and indicates that paired helper NLRs function through distinct mechanisms. However, whereas we now have a clearer understanding of the beginning and final states of NRC activation, the precise dynamics of homodimer conversion into hexamers remains unanswered. Importantly, how sensor NLRs trigger this process without becoming stably integrated into the helper oligomers is a key question which remains technically challenging to address due to the transient nature of sensor-NRC interactions (33). Another interesting question pertains to the contribution of lipids and the plasma membrane to resistosome assembly. NRCs and other CC-NLRs have been shown to shift from cytoplasm to plasma membrane-associated puncta upon activation (17, 31). Whether resistosomes assemble in the cytoplasm and then insert into membranes, or whether primed intermediates first accumulate at the membrane and then oligomerize by the aid of phospholipids remains to be determined. Obtaining structures of additional NLRs in different states of activation in their membrane context will hopefully shed light on the precise contributions of membranes to immune receptor activation. In this context, AlphaFold 3 will surely contribute to tackling these questions and complementing experimental data.

In conclusion, while the pentameric AtZAR1 resistosome kicked-off a golden age for plant NLR biology, our understanding of NLR activation is moving beyond the AtZAR1 model. Further structures of NLRs covering a broader range of plant phylogeny will surely uncover more structural and functional diversity than previously anticipated, allowing us to advance our understanding of plant NLR activation beyond singleton NLRs.

## Materials and methods

### Plant growth conditions

*nrc2/3/4* CRISPR mutant *N. benthamiana* lines were grown in a controlled environment growth chamber with a temperature range of 22 to 25 °C, humidity of 45% to 65% and a 16/8-hour light/dark cycle.

### Plasmid constructions

The Golden Gate Modular Cloning (MoClo) kit (34) and the MoClo plant parts kit (35) were used for cloning, and all vectors are from this kit unless specified otherwise. PVX CP-eGFP, Rx-6xHA and NbNRC2^EEE^-3xFLAG used in protein purification were all previously reported (17). NbNRC2 and NbNRC2 hexamerization mutants used in **Fig. 2** were cloned into the binary vector pJK001c, with a 2×35S promoter (pICSL51288), 35S terminator (pICSL41414) and C-terminal 3xFLAG tag (pICSL5007) (36). Cloning design and sequence analysis were done using Geneious Prime (v2021.2.2; https://www.geneious.com).

### Transient protein expression in *N. benthamiana* by agroinfiltration

Effectors and NLR immune receptors of interest were transiently expressed according to previously described methods (37). Briefly, leaves from 4–5-week-old plants were infiltrated with suspensions of *A. tumefaciens* GV3101 pM90 strains transformed with expression vectors coding for different proteins indicated. Final OD_600_ of all *A. tumefaciens* suspensions were adjusted in infiltration buffer (10 mM MES, 10 mM MgCl_2,_ and 150 µM acetosyringone (pH 5.6)). Final OD_600_ used was 0.3 for all NRC2 variants used, 0.1 for Rx and 0.1 for PVX CP, adding up to a total OD_600_ of 0.5.

### Protein purification from *N. benthamiana*

Around 30 leaves of *nrc2/3/4* KO *N. benthamiana* were agroinfiltrated as described above to transiently express NbNRC2^EEE^-3xFLAG, Rx-V5 and PVX CP-eGFP. Tissue was harvested after 3 days and snap-frozen in liquid nitrogen. All samples were stored at −80° C until the protein was finally extracted. The entire purification process was completed in the same day for each preparation. On the day of purification, the frozen tissue was ground into fine powder in a mortar and pestle that was pre-cooled with liquid nitrogen. 20 g of ground powder were resuspended with extraction buffer (100 mM Tris-HCl pH 7.5, 150 mM NaCl, 1 mM MgCl_2_ 1 mM EDTA, 10% glycerol, 10 mM DTT, 1x cOmplete EDTA-free protease inhibitor tablets (Sigma) and 0.2% v/v IGEPAL). 20 g of ground tissue was resuspended in ice-cold extraction buffer at a 1:4 w/v ratio (20 g of powder in 80 mL of extraction buffer). After vortexing and resuspending the powder in this buffer, the crude extract was spun down for 10 minutes at max. speed. The supernatant was transferred to a new tube and centrifuged again for 10 minutes at max. speed. This second supernatant was filtered using Miracloth (Merck). 200 ml of anti-FLAG beads were added to the supernatant and incubated at 4° C for 90 minutes. The tubes were in constant rotation to prevent the beads from sedimenting. The protein-bound beads were collected on an open column and washed with 10 ml of wash buffer (100 mM Tris-HCl pH 7.5, 150 mM NaCl, 1 mM MgCl_2_ 1 mM EDTA, 10% glycerol and 0.2% v/v IGEPAL). The washed beads were collected in an Eppendorf tube and eluted with the final isolation buffer in 200 μL volume (100mM Tris HCl (pH 7.5), 150 mM NaCl, 1 mM MgCl_2_, 3% glycerol) supplemented with 0.5 mg/ml 3xFLAG peptide. The eluted protein was analyzed on the SDS-PAGE to assess sample quality and purity. About 2.4 mg/ml concentration was obtained for NRC2^EEE^-3xFLAG from each purification, as determined by absorption at 280 nm. FLAG-eluted pure protein samples were used for cryo-EM studies.

### Negative staining

A 3.5 μL of purified NRC2^EEE^ sample, diluted to 0.3 mg/mL was applied to 400 mesh copper grids with continuous carbon (Agar Scientific) that was glow-discharged using a PELCO Easiglow (Ted Pella) for 30 seconds at 8 mA. After 30 seconds, the sample was blotted using Sartorius 292 filter paper and immediately washed with two consecutive drops of 100 μL distilled water each. After washing, 3.5 μL of 2% (w/v) uranyl acetate stain in H_2_O was applied and after 30 seconds, the excess stain was blotted off and air-dried. Grids were examined in a FEI Talos F200C transmission electron microscope operated at 200 keV, equipped with a Falcon 4i direct electron detector (Thermo Fisher Scientific). Fifty representative micrographs were recorded using EPU v3.4.0.5704 (Thermo Fisher Scientific) with a total dose of 20 e-/Å2, nominal magnification of 73 kX, and calculated pixel size of 1.7 Å. Particles were auto-picked using Cryo-SPARC (38) without any template. 2D-classification of the auto-picked particles (∼2500 particles) from these images revealed particles resembling the putative NRC2 hexamers.

### Cryo-EM sample preparation and data collection

Quantifoil R 2/1 on copper 300 mesh grids were used for Cryo-EM grid preparation. A 3.5 μL of 2.4 mg/ml of NRC2^EEE^ sample was applied over negatively glow discharged Quantifoil R 2/1 grids coated with graphene oxide. The sample was applied inside the chamber of a Vitrobot Mark IV (Thermo Fisher Scientific) at 4 °C and 90% humidity and vitrified in liquid ethane. Without graphene oxide coating, no particles were observed in vitreous ice, so graphene-coated grids were used. The Cryo-EM images were collected on a FEI Titan Krios (Thermo Fisher Scientific) operated at 300 keV equipped with a K3 direct electron detector (Gatan) after inserting an energy filter with a slit width of 20 eV. Micrographs were collected with a total dose of 50 e^−^/Å^2^, nominal magnification of 105 kx, giving a magnified pixel size of 0.828 Å. Images were collected as movies of 50 fractions with defocus values ranging from −1.5 µm to −2.7 µm with 2 exposures per hole. A total of 6135 movies were collected for the image processing and 3D reconstruction using CryoSPARC (38, 39).

The micrograph movies were imported into CryoSPARC and subjected to drift correction using MotionCor2 (40) and CTFFIND4.0 was used for fitting of contrast transfer function and defocus estimation. The Laplacian-of-Gaussian auto-picker in CryoSPARC was used for automatic reference-free particle picking (38, 39). The particles were extracted in 256-pixel box and subjected to several rounds of 2D classification with a circular mask of 180 Å. The 2D classes revealed a clear hexamer with secondary structural features in different views. Clean 2D-classes were selected (664,305 particles) and were subjected to 3D classification using an *ab initio* 3D model generated by Cryo-SPARC (38). The best 3D class revealing the protein fold consisted of 229,347 particles and was subjected to 3D refinement using Refine3D using that 3D class as a reference with a circular mask of 180 Å. Following examination of the C1 refined map, a C2 symmetry was applied. After particle polishing, CTF refinement and postprocessing, the final average resolution was 2.9 Å as estimated by the Fourier shell correlation (FSC = 0.143). The local resolution plot was calculated using PHENIX 3.0 (41). The final map revealed distinct domain boundaries of the NRC2 protein fold with clear secondary structural features, helical twists and individual beta strands and allowed us to place each NRC2 protomer and initiate model building at this resolution. A significant number of particle views are down the short dimension, owing to graphene oxide backing. The final map revealed the CC, NB-ARC and LRR domains of the NRC2 molecule.

### Model building and refinement

An initial models of the NRC2 monomer was generated with AlphaFold2 (29) and fitted approximately within the density by structural alignment to a model of the entire NRC2 hexamer built using ModelAngelo (42). Iterations of manual adjustment in Coot, de novo model building and real-space refinement in Phenix were performed with the consensus EM map. (41, 43). This model was validated using PHENIX and MolProbity (41, 44). Figures were made using ChimeraX (45). The interface residues and the buried surface area were evaluated using PyMOL and the PISA server (46, 47). A distance cut-off 5 Å is used to define residues at the interface to accommodate all short- and long-range interacting residues. Further details on Cryo-EM data processing statistics can be found in **Table S1**.

### Extraction of total proteins for BN-PAGE and SDS-PAGE assays

Four to five-week-old plants were agroinfiltrated as described above with constructs of interest and leaf tissue was collected 2 days post agroinfiltration in experiments. Final OD_600_ used was 0.3 for 0.3 for all NRC2-WT or variants used, 0.1 for Rx and 0.1 for PVX CP, and 0.1 for GFP and a total OD_600_ of 0.5. BN-PAGE was performed using the Bis-Tris Native PAGE system (Invitrogen) according to the manufacturer’s instructions. Leaf tissue was ground using a Geno/Grinder tissue homogenizer and total protein was subsequently extracted and homogenized extraction buffer. For NRC2, GTMN extraction buffer was used (10% glycerol, 50 mM Tris-HCl (pH 7.5), 5 mM MgCl_2_ and 50 mM NaCl) supplemented with 10 mM DTT, 1x protease inhibitor cocktail (SIGMA) and 0.2% Nonidet P-40 Substitute (SIGMA). Samples were incubated in extraction buffer on ice for 15 minutes with short vortex mixing at every 2 minutes. Following incubation, samples were centrifuged at 5,000 x*g* for 15 minutes and the supernatant was used for BN-PAGE and SDS-PAGE assays.

### BN-PAGE assays

For BN-PAGE, samples extracted as detailed above were diluted as per the manufacturer’s instructions by adding NativePAGE 5% G-250 sample additive, 4x Sample Buffer and water. After dilution, samples were loaded and run on Native PAGE 3%-12% Bis-Tris gels alongside either NativeMark unstained protein standard (Invitrogen) or SERVA Native Marker (SERVA). The proteins were then transferred to polyvinylidene difluoride membranes using NuPAGE Transfer Buffer using a Trans-Blot Turbo Transfer System (Bio-Rad) as per the manufacturer’s instructions. Proteins were fixed to the membranes by incubating with 8% acetic acid for 15 minutes, washed with water and left to dry. Membranes were subsequently re-activated with methanol in order to correctly visualize the unstained native protein marker. Membranes were immunoblotted as described below.

### SDS-PAGE assays

For SDS-PAGE, samples were diluted in SDS loading dye and denatured at 72 °C for 10 minutes. Denatured samples were spun down at 5,000 x*g* for 3 minutes and supernatant was run on 4%-20% Bio-Rad 4%-20% Mini-PROTEAN TGX gels alongside a PageRuler Plus prestained protein ladder (Thermo Scientific). The proteins were then transferred to polyvinylidene difluoride membranes using Trans-Blot Turbo Transfer Buffer using a Trans-Blot Turbo Transfer System (Bio-Rad) as per the manufacturer’s instructions. Membranes were immunoblotted as described below.

### Immunoblotting and detection of BN-PAGE and SDS-PAGE assays

Blotted membranes were blocked with 5% milk in Tris-buffered saline plus 0.01% Tween 20 (TBS-T) for an hour at room temperature and subsequently incubated with desired antibodies at 4 °C overnight. Antibodies used were anti-GFP (B-2) HRP (Santa Cruz Biotechnology), anti-Myc (9E10) HRP (Roche), and anti-FLAG (M2) HRP (Sigma), all used in a 1:5000 dilution in 5% milk in TBS-T. To visualize proteins, we used Pierce ECL Western (32106, Thermo Fisher Scientific), supplementing with up to 50% SuperSignal West Femto Maximum Sensitivity Substrate (34095, Thermo Fishes Scientific) when necessary. Membrane imaging was carried out with an ImageQuant LAS 4000 or an ImageQuant 800 luminescent imager (GE Healthcare Life Sciences, Piscataway, NJ). Rubisco loading control was stained using Ponceau S (Sigma) or Ponceau 4R (AG Barr).

### Structural modeling and processing

ColabFold v1.5.5 was used to model NbNRC2a CC-NB-ARC (29, 48). The AlphaFold 3 webserver (https://golgi.sandbox.google.com/) was used to model the CC-NB-ARC domains of 11 NRC helpers, 10 CC-NLRs, 6 CC_G10_-NLRs, and 2 CC_R_-NLRs with 50 oleic acids as a proxy for the plasma membrane (**Table S2**) (26). The default seed was set to 1. NbNRC4c and NbNRG1 hexamers were modelled with seed 11. To replicate NbNRC2a oligomers, we generated 9 additional models with randomized seed numbers, in addition to the default seed (**Table S2**). From each modeling run, only the top-ranked model (model_0) was retained and processed. The ipTM, pTM, chain-pair ipTM, and minimum chain pTM values were extracted from the AlphaFold 3 JSON files containing model metadata using custom scripts and plotted in R. The AlphaFold 3 models matching the cryo-EM structures of NbNRC2a (PDB), AtZAR1 (6j5t), and TmSr35 (7xe0) were aligned to the cryo-EM structures using the matchmaker function of ChimeraX with default options (45). The contact points from AlphaFold structures were extracted using the ChimeraX [alphafold contacts #1/a to #1/b distance 4.5] (45). All structures were visualized using ChimeraX and assembled manually (45). All scripts are available at [github.com/amiralito/NRC2Hexamer]. AlphaFold predicted structures are deposited on Zenodo (49).

## Acknowledgements

We thank Doryen Bubeck (Imperial College London), and Raoul Frijters (Rijk Zwaan) for useful comments and suggestions. We thank N. Lukoyanova (Birkbeck, University of London, London, UK) for support with Cryo-EM imaging. M.P.C. thanks C. Rica and B. Variegatus for inspiration during the writing process. Cryo-EM data for this investigation were collected at ISMB EM facility, which is supported by the Wellcome Trust (202679/Z/16/Z and 206166/Z/17/Z). We thank all members of the TSL Support Services for their invaluable assistance.

## Funding

The authors received funding from the sources listed below. The funders had no role in the study design, data collection and analysis, decision to publish, or preparation of the manuscript. The Gatsby charitable foundation, Biotechnology and Biological Sciences Research Council (BBSRC) BB/P012574 (Plant Health ISP), BBSRC BBS/E/J/000PR9795, BBSRC BBS/E/J/000PR9796 (Plant Health ISP – Response), BBSRC BBS/E/J/000PR9797 (Plant Health ISP – Susceptibility), BBSRC BBS/E/J/000PR9798 (Plant Health ISP – Evolution), BBSRC BB/V002937/1, BBSRC BB/T006102/1, BBSRC BB/X01102X/1, European Research Council (ERC) 743165 and UK Research and Innovation (UKRI) Future Leaders Fellowship MR/X033481/1.

## Author contributions

Conceptualisation: J.M., M.P.C., T.O.B., M.W., S.K.

Methodology: J.M., A.T., A.P., J.R., M.W.

Data curation: J.M., A.T., J.R., M.W.

Formal analysis: J.M., A.T., M.W.

Investigation: J.M., A.T., J.R., M.W. Resources: S.K., J.K.

Data curation: J.M., A.T., J.R, M.W.

Writing – original draft: J.M., A.T., M.P.C, M.W., S.K.

Writing – review and editing: J.M., A.T., M.P.C., A.P., J.R., J.K., T.O.B., M.W., S.K.

Visualisation: J.M., A.T., M.W.

Supervision: J.M., M.P.C., M.W., S.K.

Project Administration: J.M., M.P.C., A.T., M.W., S.K.

Funding acquisition: M.W., S.K.

## Competing interest statement

T.O.B., and S.K. receive funding from industry on NLR biology and cofounded a start-up company (Resurrect Bio Ltd.) on resurrecting disease resistance. M.P.C., J.K. and S.K. have filed patents on NLR biology. M.P.C. and J.K. have received fees from Resurrect Bio Ltd.

## Supplementary information

**Fig. S1:**
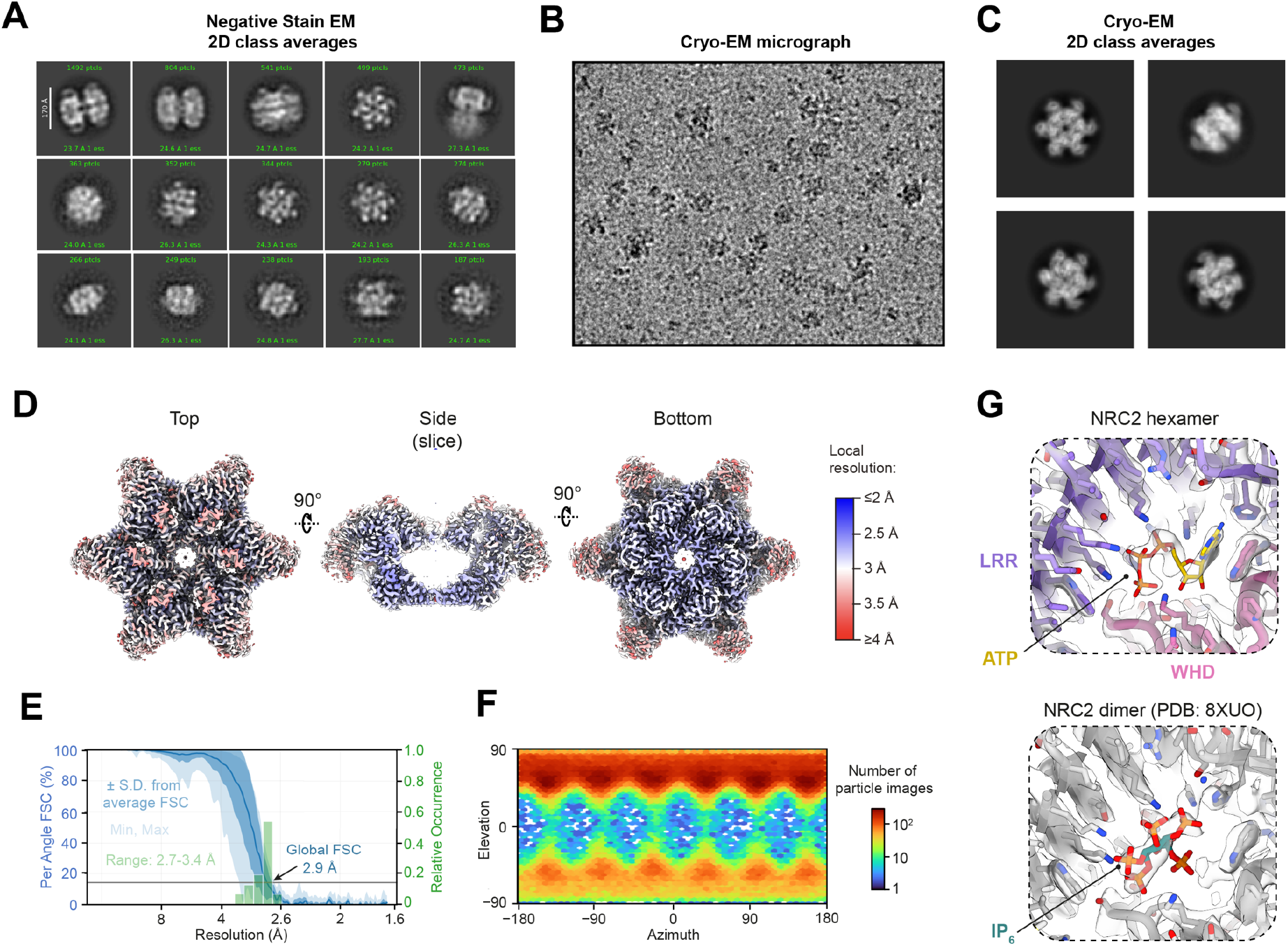
Cryo-EM analysis reveals homohexameric resistosome structure of NbNRC2 activated with Rx and *Potato virus X* coat protein. (**A**) Analysis of purified NbNRC2^EEE^ by single-particle negative-stain electron microscopy and the longest dimension in the 3D reconstruction is 17 nm as indicated. (**B**) Representative cryo-EM micrograph of NbNRC2^EEE^. (**C**) Selected cryo-EM 2D class averages from the final particle set. (**D**) Local resolution estimate for consensus NbNRC2^EEE^ resistosome after reconstruction. (**E**) Fourier shell correlation plot for consensus NbNRC2^EEE^ resistosome reconstruction. (**F**) Angular distribution plot for consensus NbNRC2^EEE^resistosome reconstruction. (**G**) Cryo-EM reconstruction of the NRC2 hexamer (top) shows additional density in the concave surface of the LRR repeat consistent with a nucleotide triphosphate. The corresponding position in NRC2 dimer reconstruction (bottom, PDB: 8XUO) contained density at the corresponding position with a different shape that was identified as IP_6_.

**Fig. S2:**
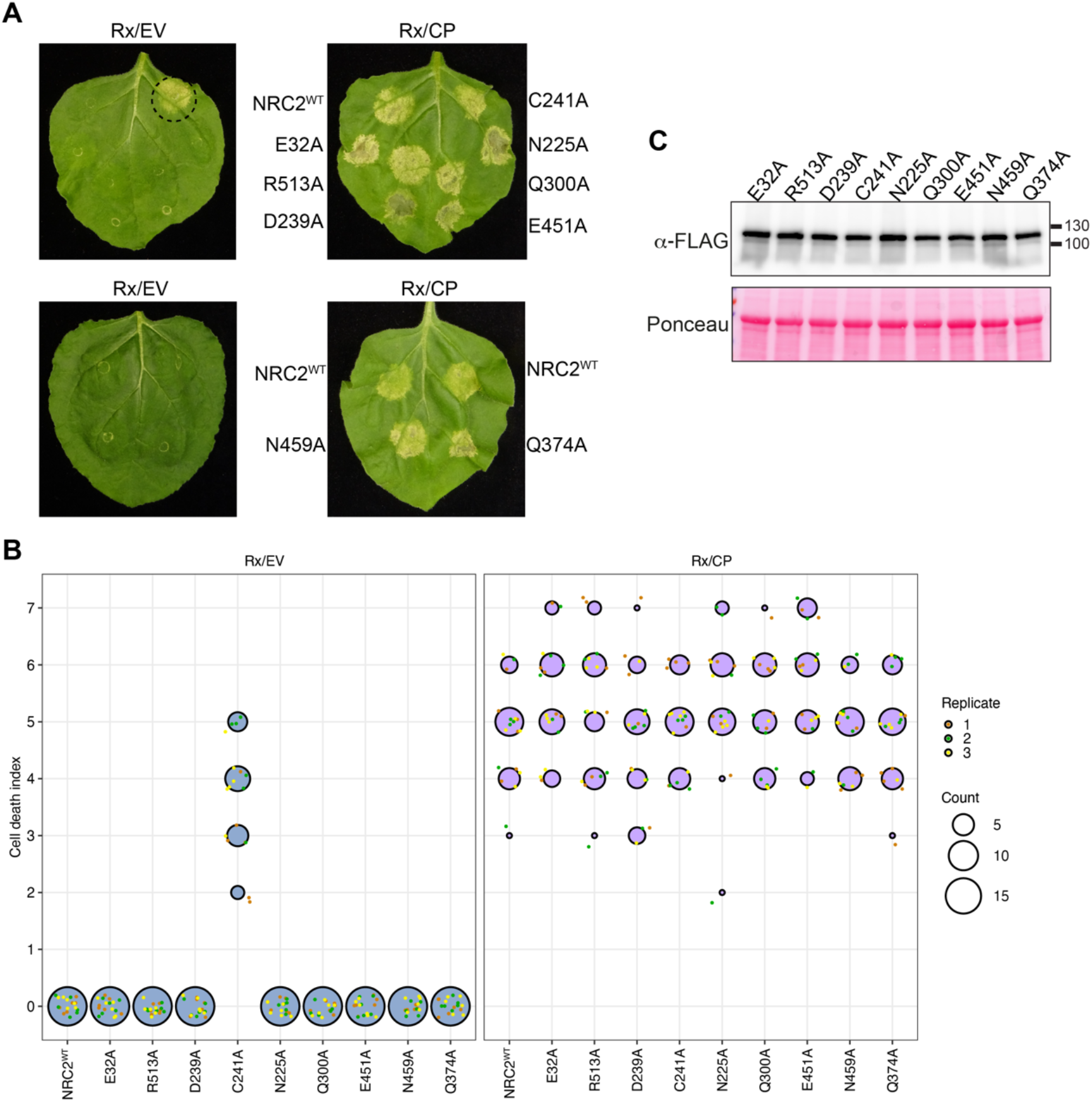
NbNRC2 single amino acid mutants in the oligomerization interface trigger cell death upon activation by Rx/CP. (**A**) Photo of representative leaves from *N. benthamiana nrc2/3/4* KO plants showing HR after co-expression of NbNRC2 variants and Rx, together with either EV or CP. **(B)** SDS-PAGE of all NbNRC2 variants tested. Total protein extracts were immunoblotted with the appropriate antisera labelled on the left. Approximate molecular weights (kDa) of the proteins are shown on the right. Rubisco loading control was carried out using Ponceau stain (PS). The experiment was repeated three times with similar results. (**C**) HR scores accompanying panel (**A**). HR was scored based on a modified 0-7 scale between 5-7 days post-infiltration (50). HR scores are presented as dot plots, where the size of each dot is proportional to the number of samples with the same score (Count). Results are based on 3 biological replicates.

**Fig. S3:**
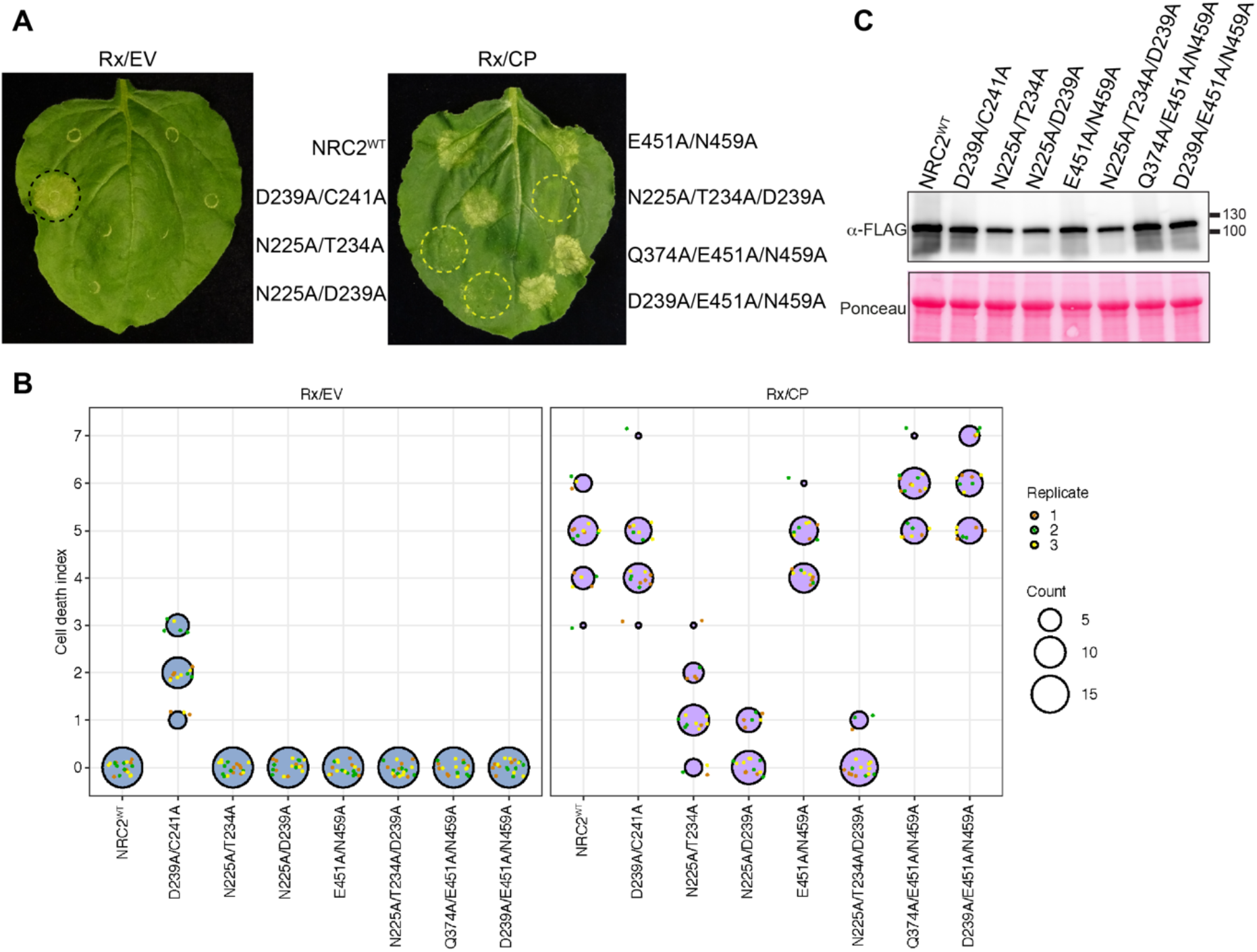
Mutations in the NB-NB inter-protomer interface abolish NbNRC2-mediated cell death. (**A**) Photo of representative leaves from *N. benthamiana nrc2/3/4* KO plants showing HR after co-expression of NbNRC2 variants and Rx, together with either EV or CP. **(B)** SDS-PAGE of all NbNRC2 variants tested. Total protein extracts were immunoblotted with the appropriate antisera labelled on the left. Approximate molecular weights (kDa) of the proteins are shown on the right. Rubisco loading control was carried out using Ponceau stain (PS). The experiment was repeated three times with similar results. (**C**) HR scores accompanying panel (**A**). HR was scored based on a modified 0-7 scale between 5-7 days post-infiltration (50). HR scores are presented as dot plots, where the size of each dot is proportional to the number of samples with the same score (Count). Results are based on 3 biological replicates.

**Fig. S4:**
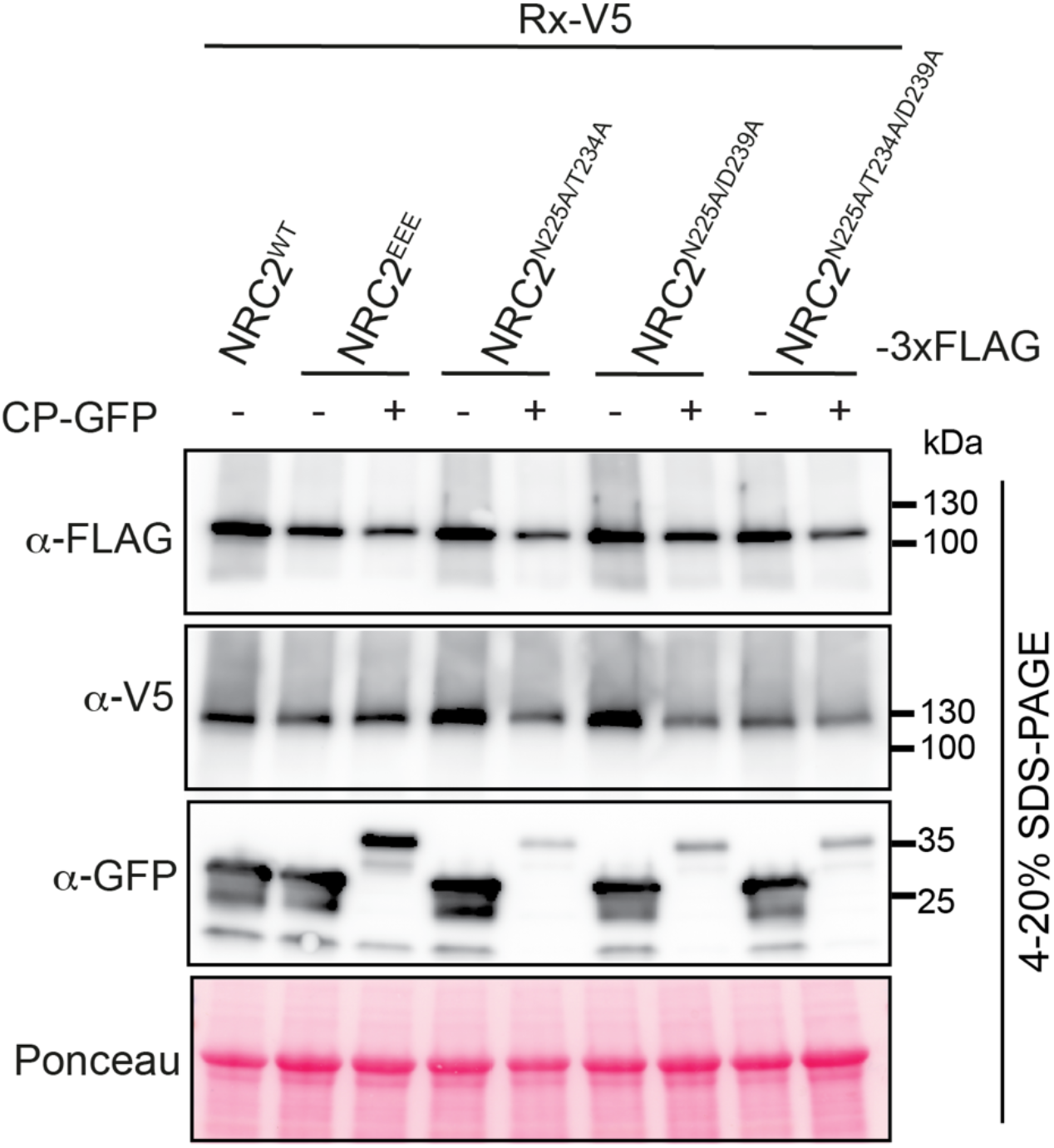
Mutants in NbNRC2 inter-protomer oligomerization interface fail to form resistosomes. SDS-PAGE of all NbNRC2 variants tested, accompanying BN-PAGE experiment in **Fig. 2**. Total protein extracts were immunoblotted with the appropriate antisera labelled on the left. Free GFP was used as a control for CP-GFP. Approximate molecular weights (in kDa) of the proteins are shown on the right. Rubisco loading control was carried out using Ponceau stain (PS). The experiment was repeated three times with similar results.

**Fig. S5.**
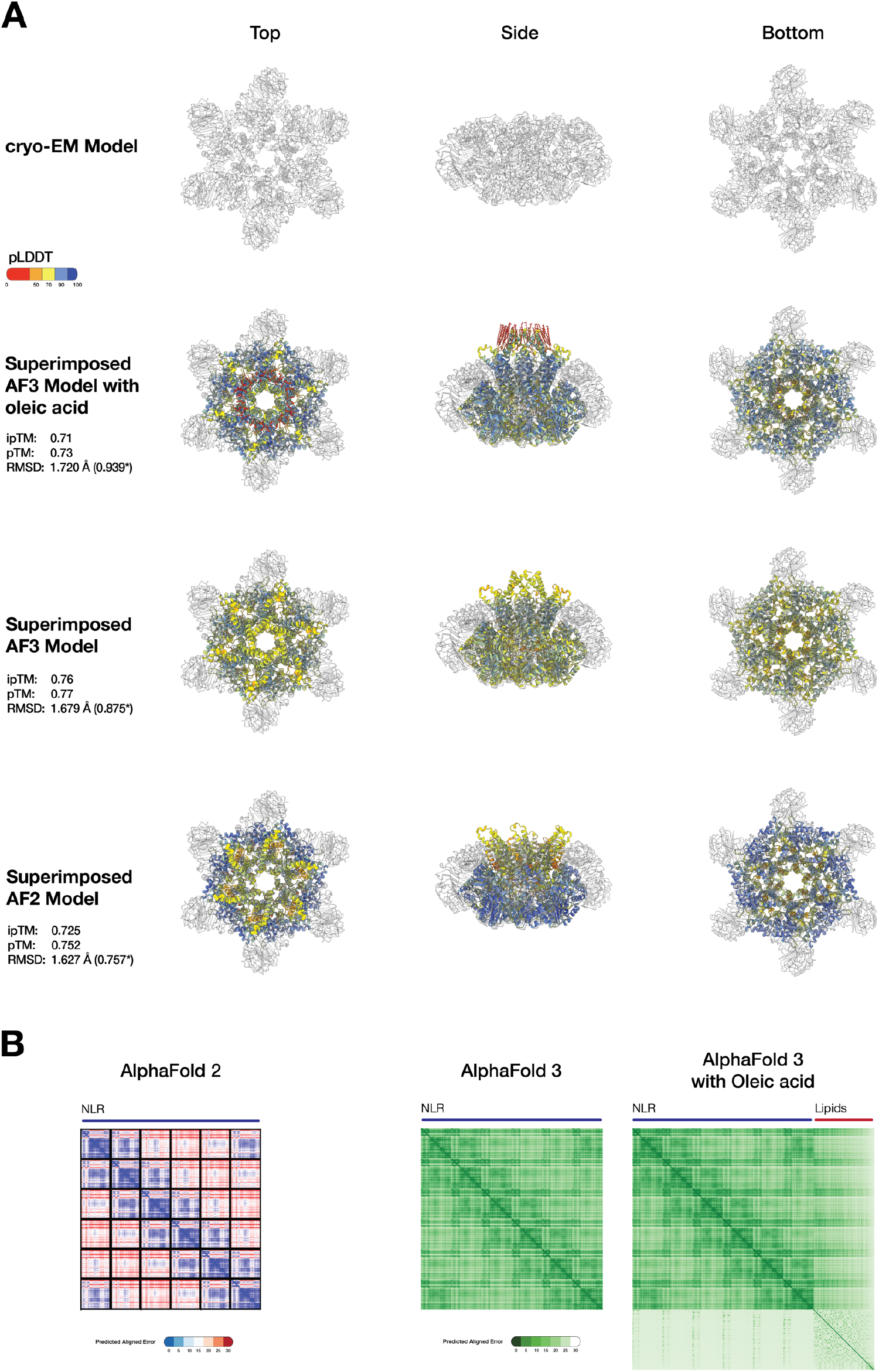
AlphaFold 3 can predict a high-confidence NbNRC2 resistosome bound to the complete funnel structure. (**A**) Superimposition of NbNRC2 predicted structures from AlphaFold 3 with and without oleic acids, and AlphaFold 2 on the cryo-EM model. AlphaFold 3 predicted structure with oleic acid had a complete and confident lipid-bound funnel structure (**B**) Predicted aligned error plots for the three models from AlphaFold 2, AlphaFold 3, and AlphaFold 3 with oleic acid.

**Fig. S6:**
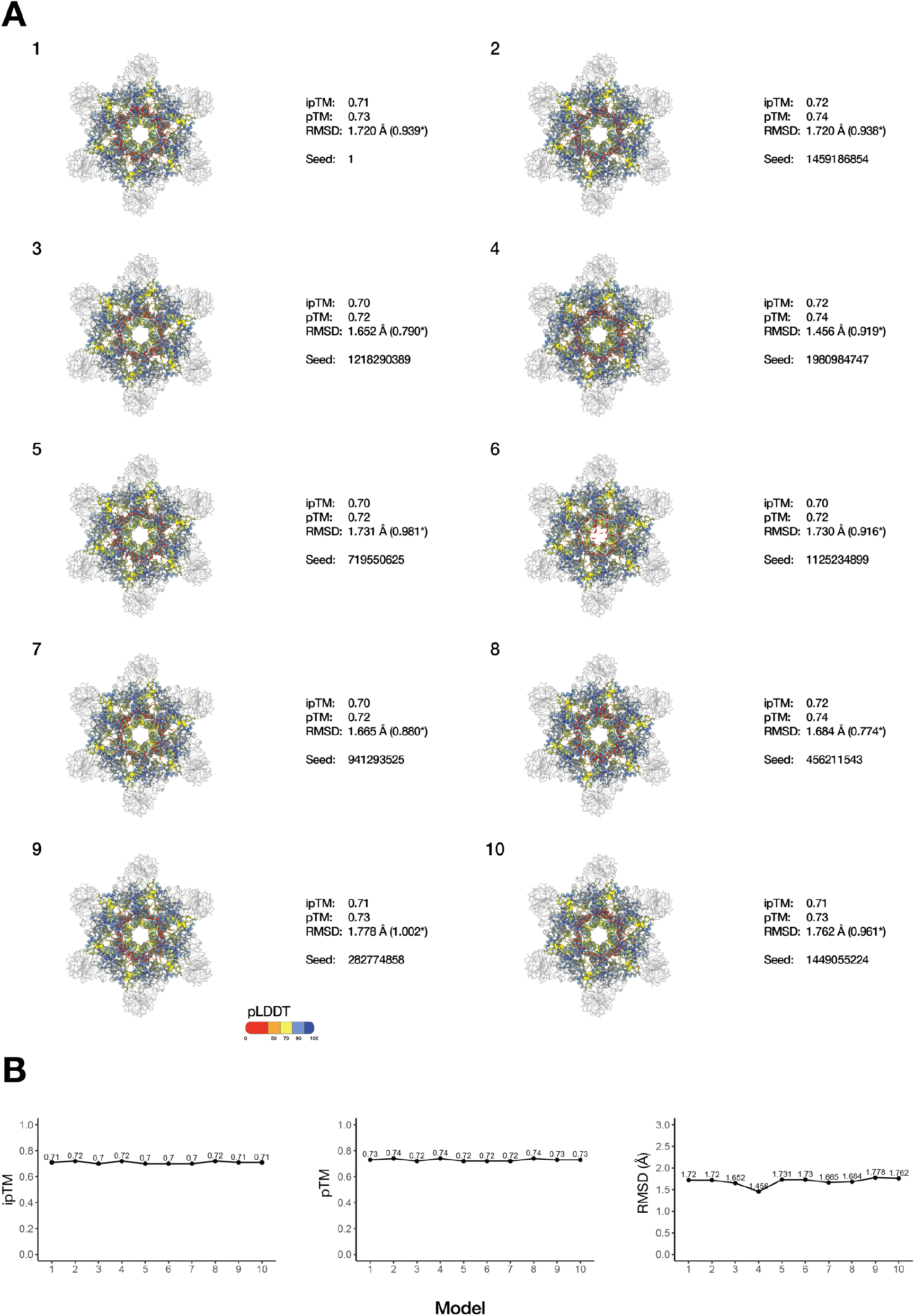
AlphaFold 3 consistently produces high-confidence NbNRC2 resistosome models. (**A**) and (**B**) 10 generated NbNRC2 resistosomes with random seeds. All models aligned with NbNRC2 cryo-EM structure with RMSDs of 1.8 Å and lower.

**Fig. S7.**
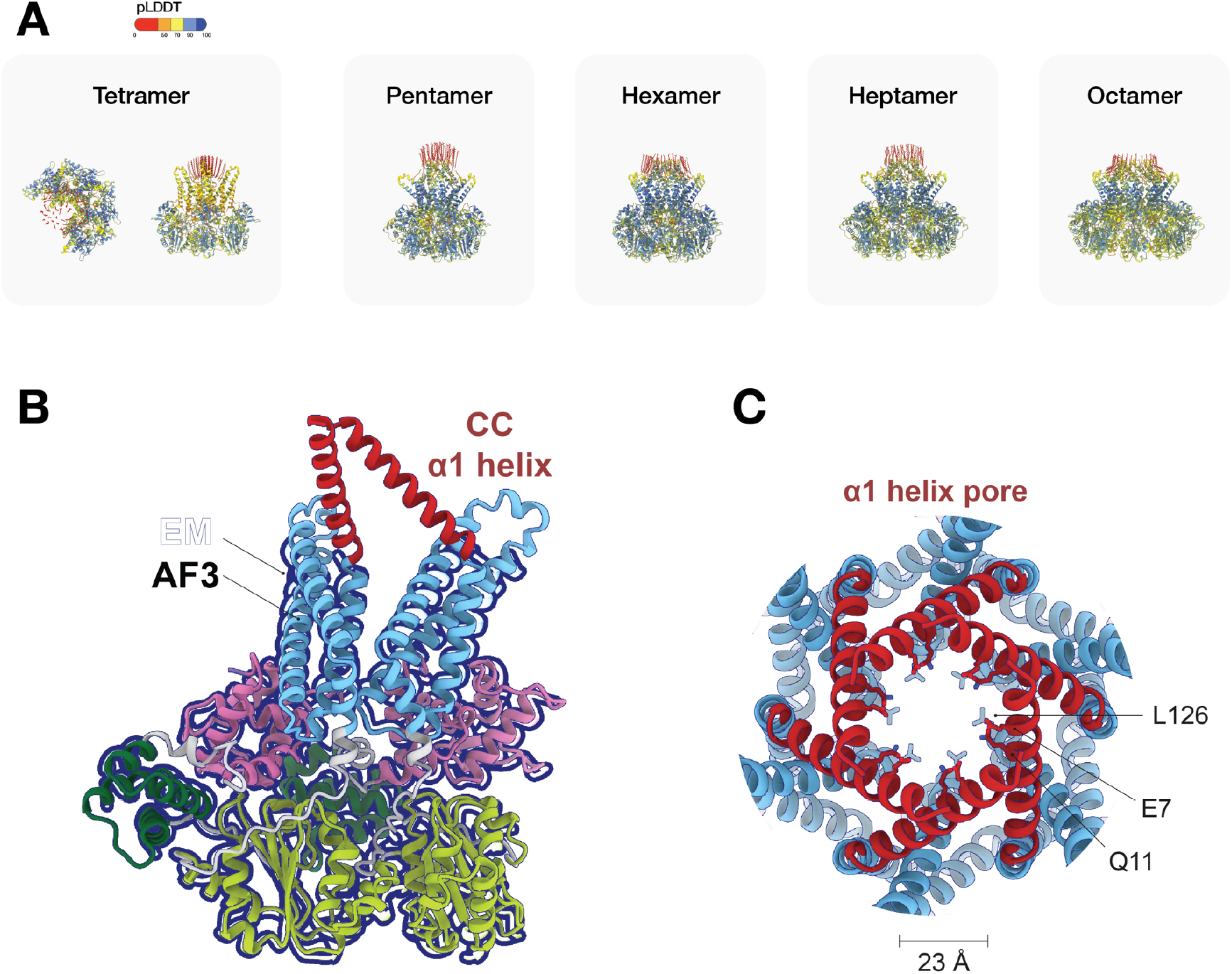
AlphaFold 3 can predict different oligomeric configurations of NbNRC2 with variable confidences. (**A**) Representative AlphaFold3 predicted models of NbNRC2 as a tetramer, pentamer, hexamer, heptamer, and octamer. (**B**) Superimposition of two adjunct protomers from AlphaFold 3 model on the cryo-EM model with relative positioning of the alpha1-helix in the CC domain. (**C**) Top view of the NbNRC2 hexamer resistosome pore with predicted alpha1-helices from AlphaFold 3.

**Fig. S8.**
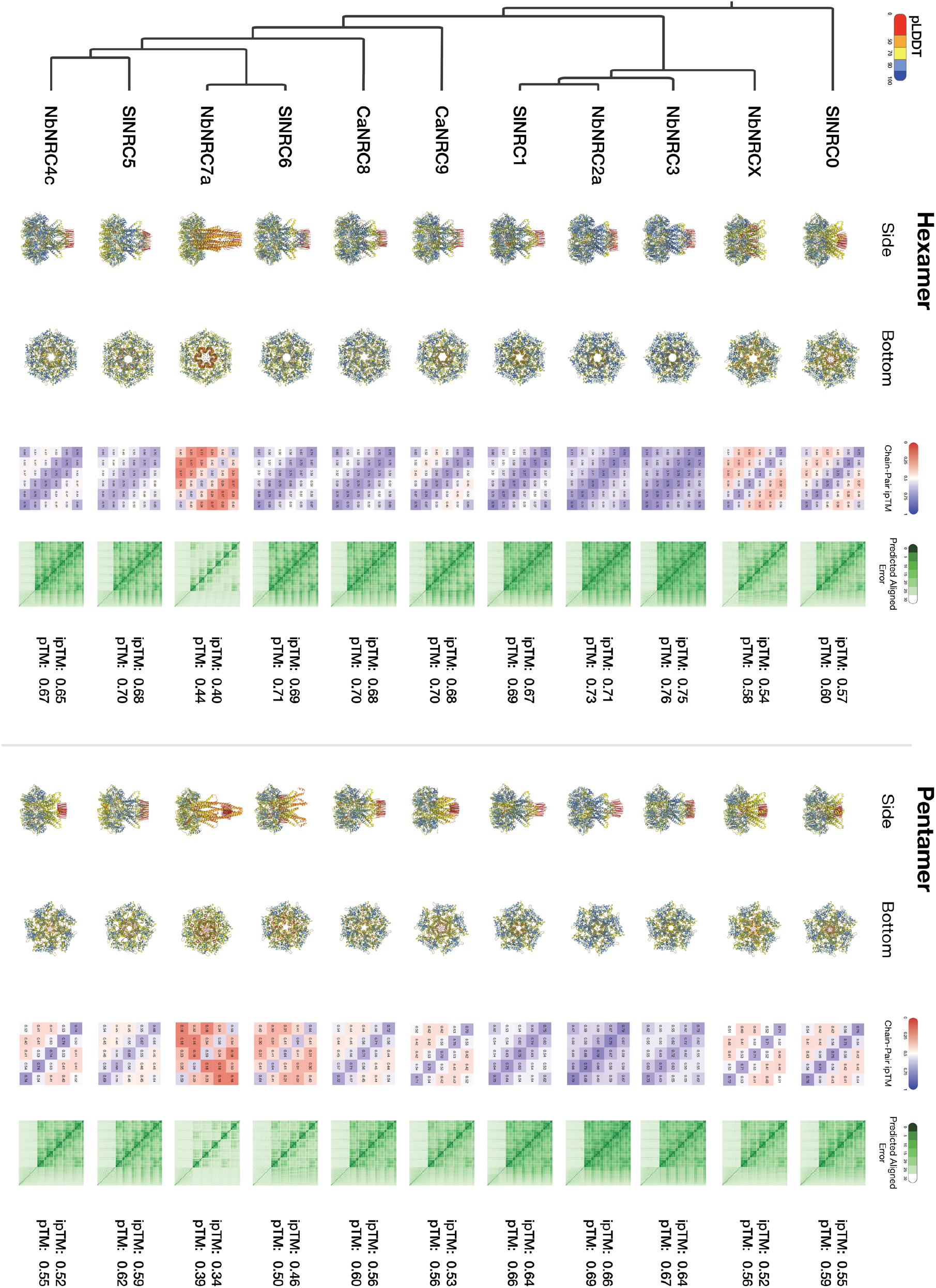
Predicted pentamer and hexamer AlphaFold 3 models from representatives from NRC helper clades. Structures are colored by pLDDT values. pTM, ipTM, Chain-Pair ipTM, and predicted aligned error metrics are provided. Phylogenetic tree adapted from Selvaraj, et al., 2023. (24).

**Fig. S9.**
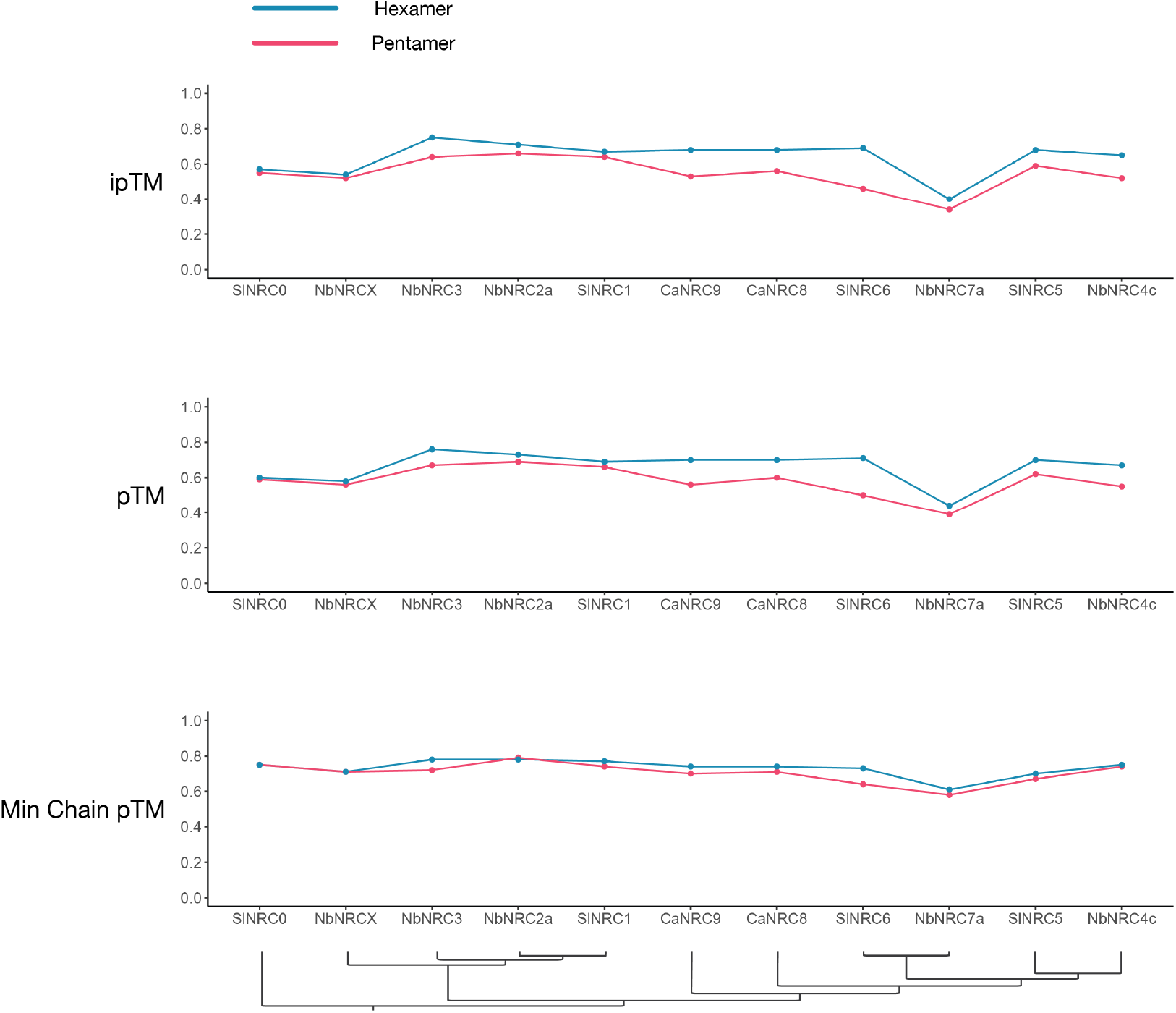
ipTM, pTM, and Minimum Chain pTM confidence metrics for modelled representatives from NRC helper clades.

**Fig. S10.**
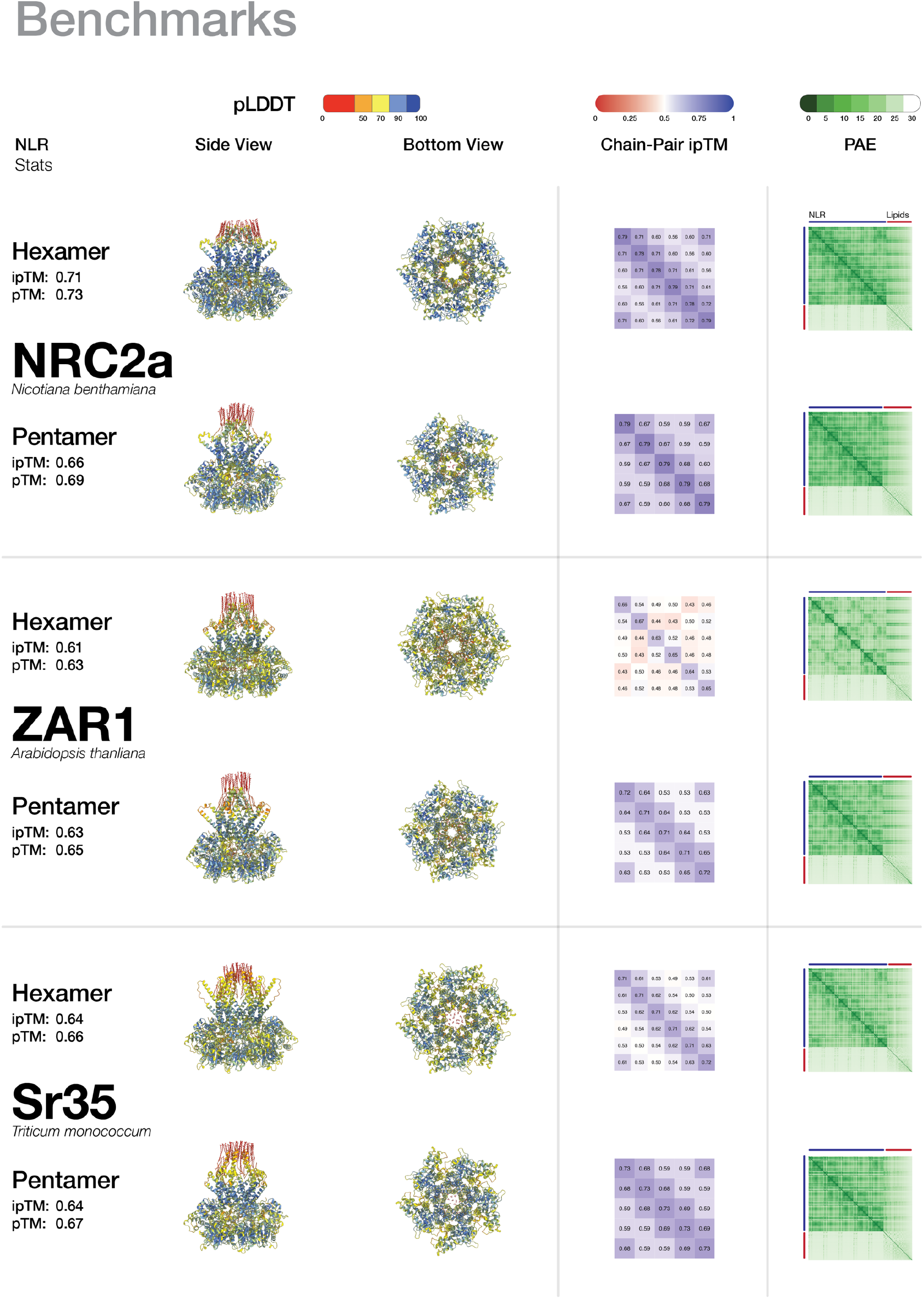
AlphaFold 3 models of NbNRC2, AtZAR1, and TmSr35 as pentamers and hexamers. Structures are colored by pLDDT values. pTM, ipTM, Chain-Pair ipTM, and predicted aligned error metrics are provided.

**Fig. S11.**
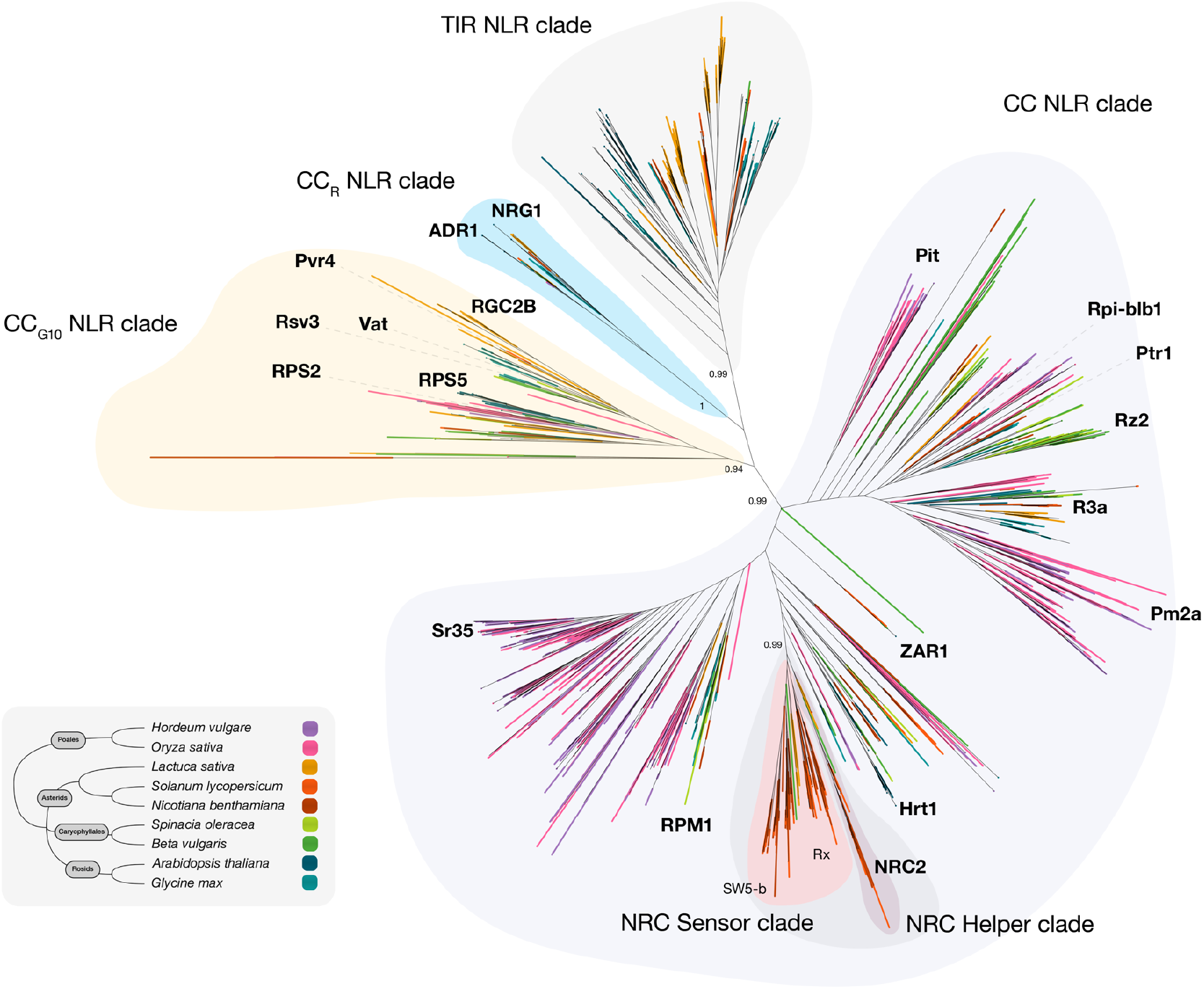
Phylogenetic tree of NLRs from nine representative species from Poales, Asterids, Caryophyllales, and Rosids. Modelled NLRs are highlighted in bold on the tree. Phylogenetic tree was adapted from Contreras, et al., 2023 (3).

**Fig. S12.**
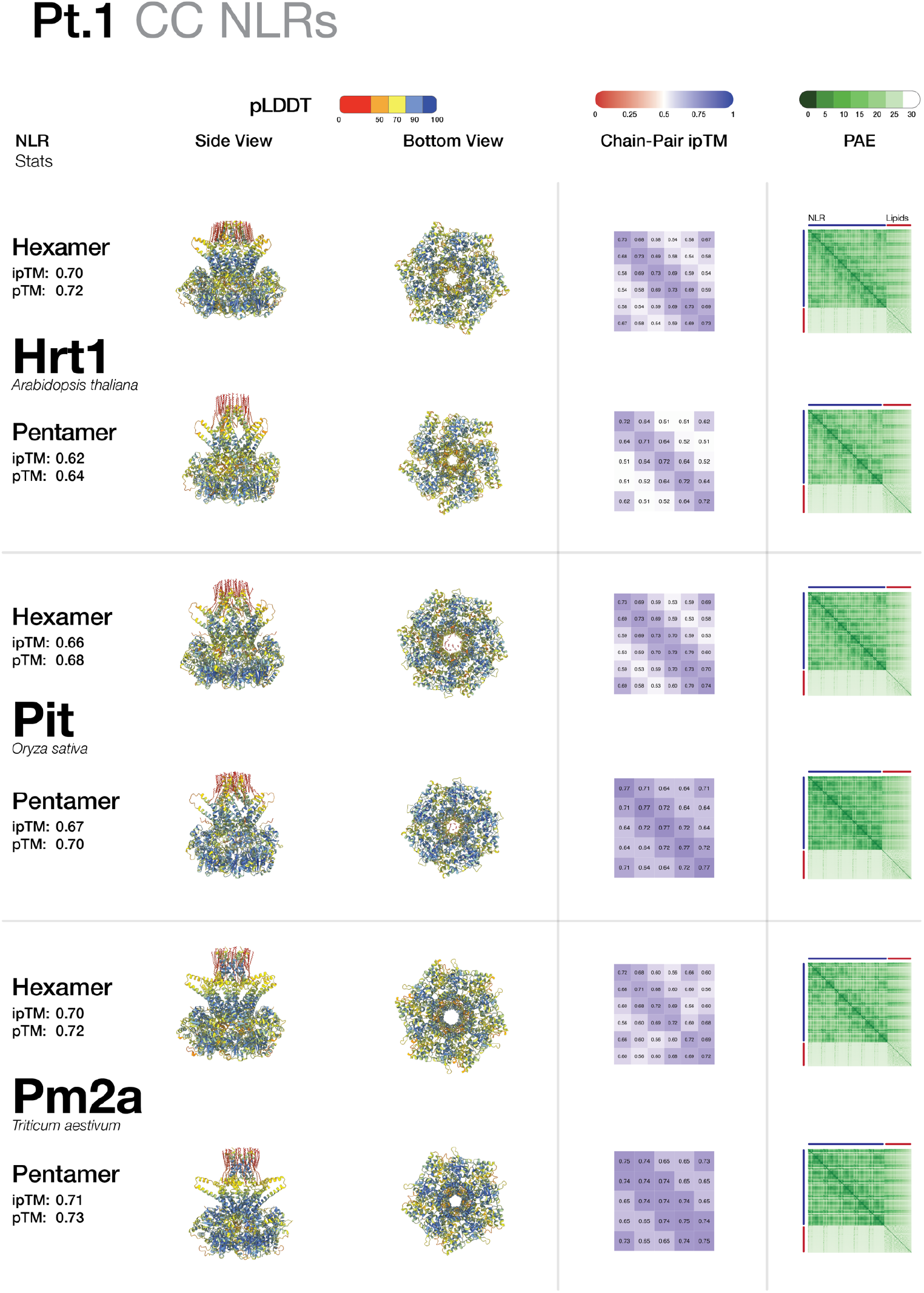
AlphaFold 3 models of selected CC-NLRs, AtHrt1, PsPit, and TaPm2a, as pentamers and hexamers. Structures are colored by pLDDT values. pTM, ipTM, Chain-Pair ipTM, and predicted aligned error metrics are provided.

**Fig. S13.**
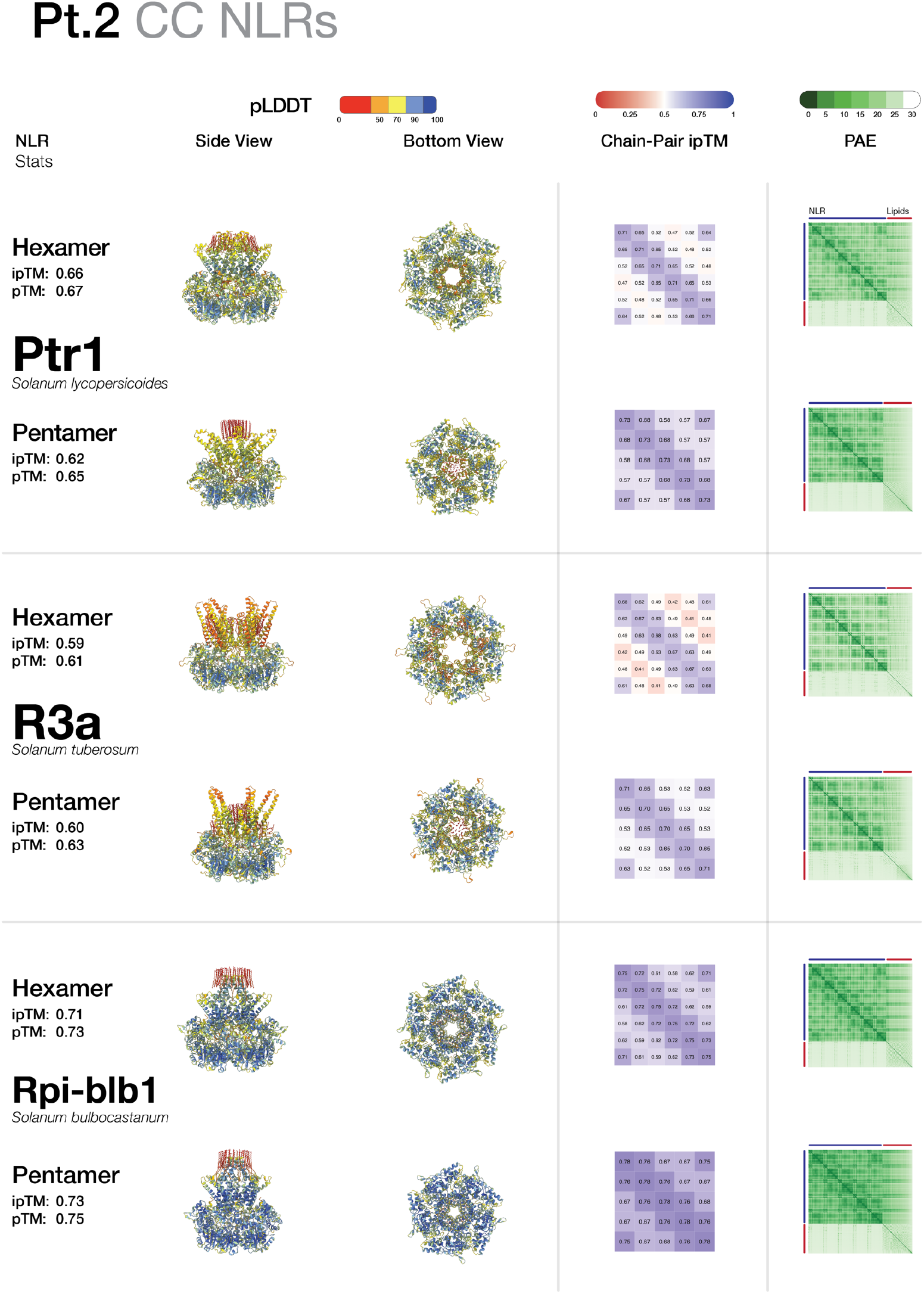
AlphaFold 3 models of selected CC-NLRs, SlPtr1, StR3a, and SbRpi-blb1, as pentamers and hexamers. Structures are colored by pLDDT values. pTM, ipTM, Chain-Pair ipTM, and predicted aligned error metrics are provided.

**Fig. S14.**
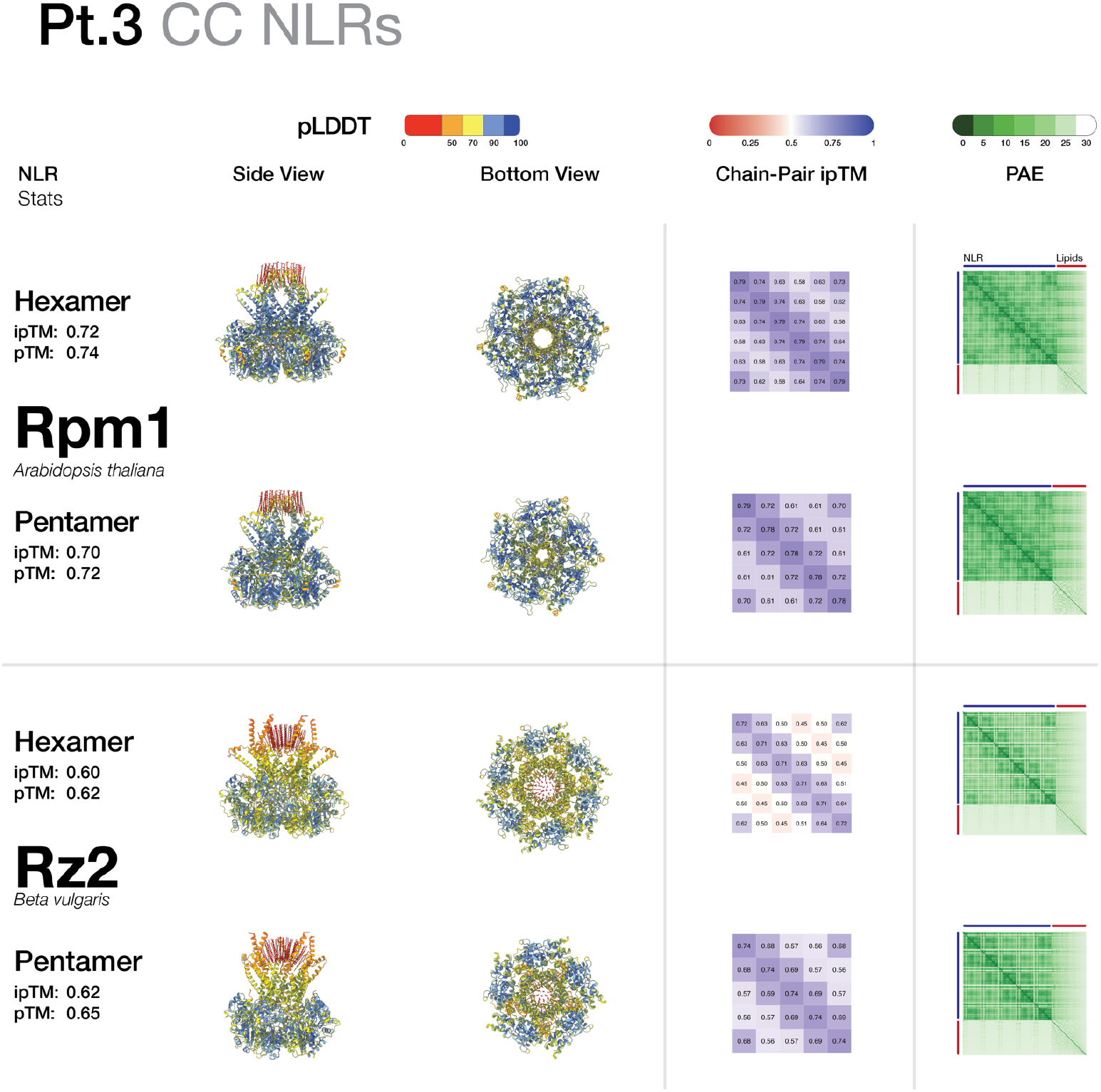
AlphaFold 3 models of selected CC-NLRs, AtRpm1, and BvRz2, as pentamers and hexamers. Structures are colored by pLDDT values. pTM, ipTM, Chain-Pair ipTM, and predicted aligned error metrics are provided.

**Fig. S15.**
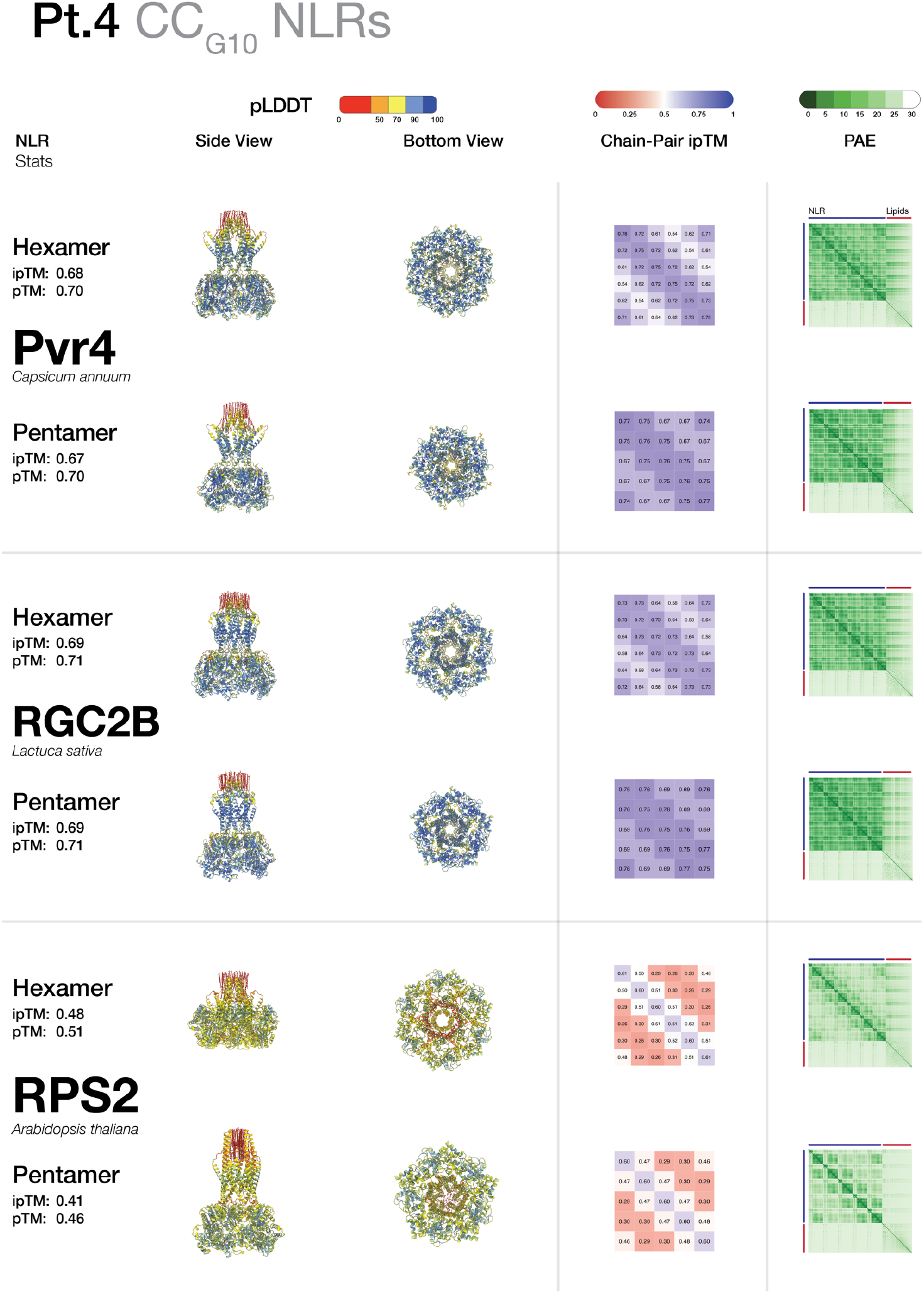
AlphaFold 3 models of selected CC_G10_-NLRs, CaPvr4, LsRGC2B, and AtRPS2, as pentamers and hexamers. Structures are colored by pLDDT values. pTM, ipTM, Chain-Pair ipTM, and predicted aligned error metrics are provided.

**Fig. S16.**
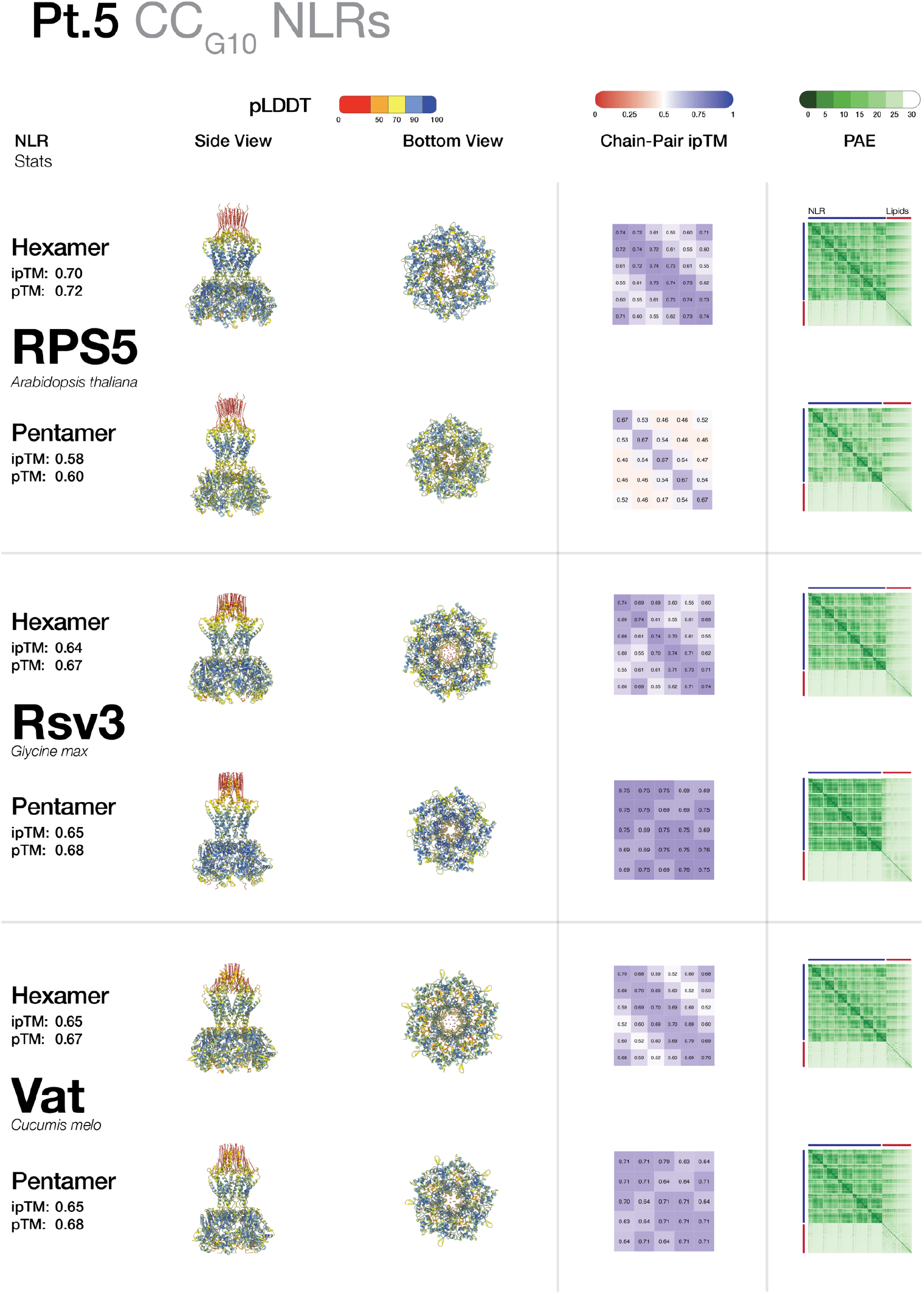
AlphaFold 3 models of selected CC_G10_-NLRs, AtRPS5, GmRsv3, and CmVat, as pentamers and hexamers. Structures are colored by pLDDT values. pTM, ipTM, Chain-Pair ipTM, and predicted aligned error metrics are provided.

**Fig. S17.**
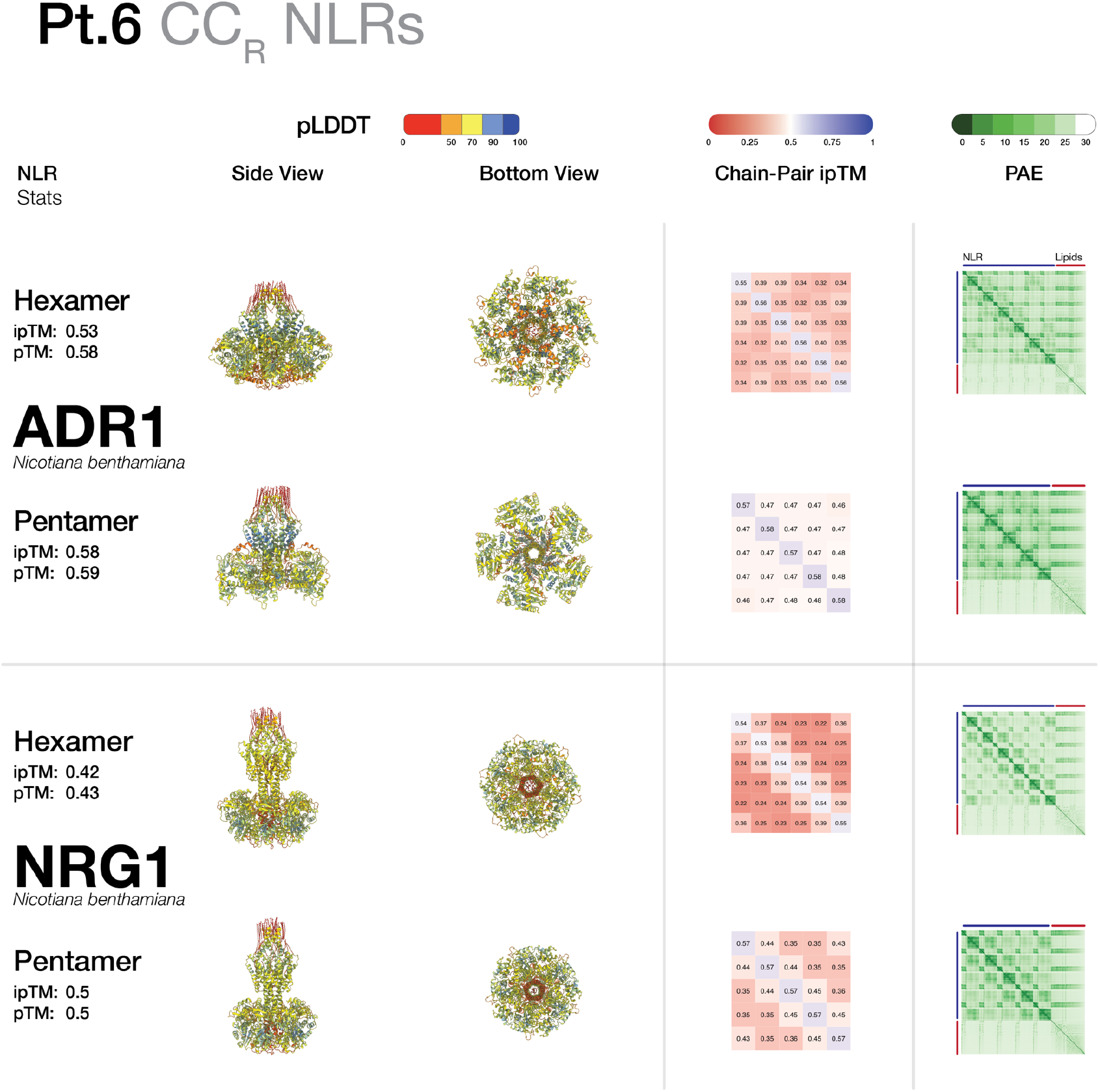
AlphaFold 3 models of selected CC_R_-NLRs, NbADR1, and NbNRG1, as pentamers and hexamers. Structures are colored by pLDDT values. pTM, ipTM, Chain-Pair ipTM, and predicted aligned error metrics are provided.

**Fig. S18:**
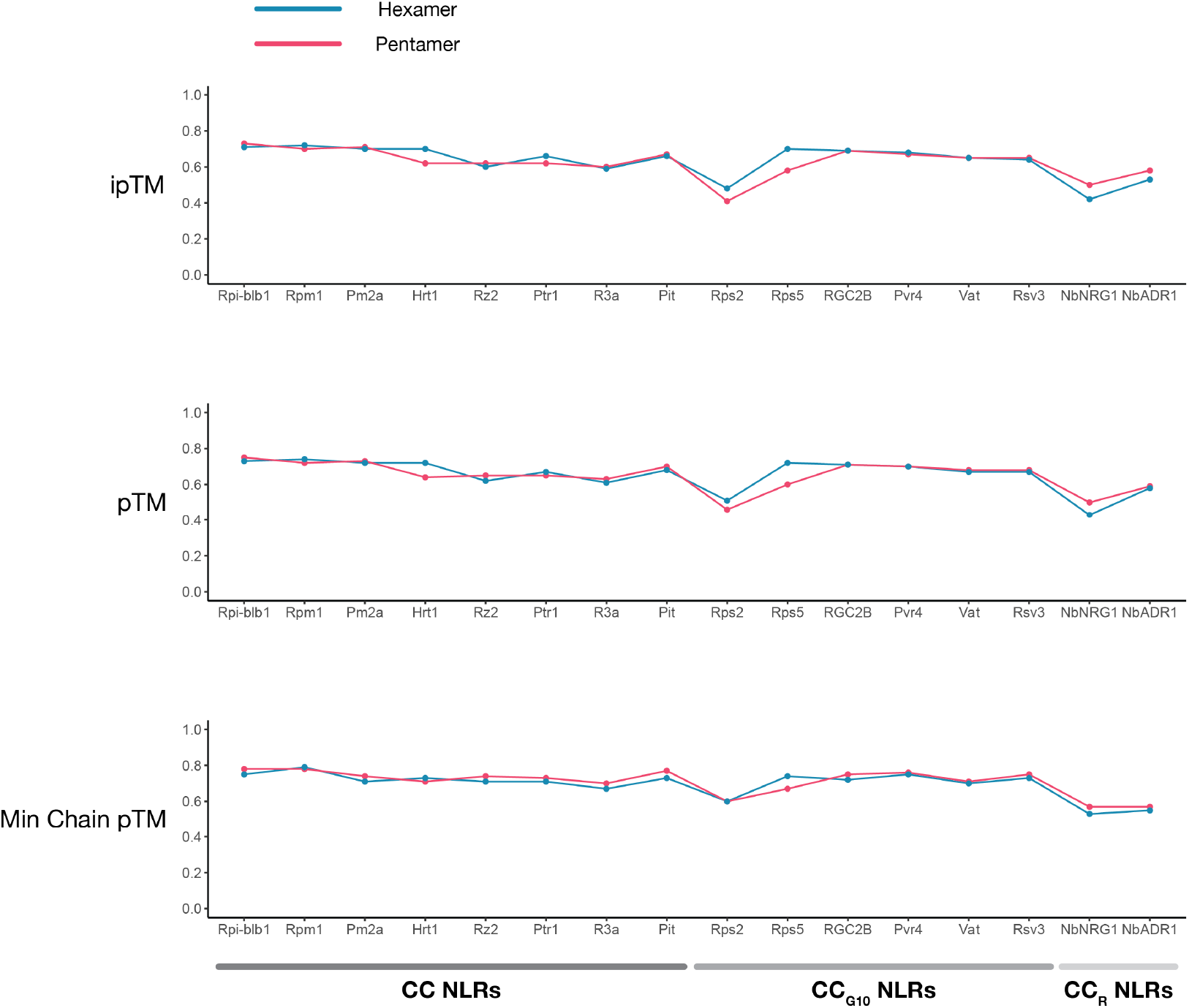
ipTM, pTM, and Minimum Chain pTM confidence metrics for modelled representative CC, CC_G10_, and CC_R_ NLRs.

**Table S1:** Cryo-EM data collection, refinement, and validation statistics.

|  |  |
| --- | --- |
| **Data collection** |  |
| Pixel size (Å) | 0.828 |
| Electron flux (e<sup>-</sup> / Å<sup>2</sup>/sec) | 25.4 |
| Micrograph exposure time (sec) | 1.95 |
| Electron fluence per frame (e<sup>-</sup> / Å<sup>2</sup>) | 49.5 |
| Number of frames | 50 |
| Total electron fluence (e<sup>-</sup> / Å<sup>2</sup>) | 25.4 |
| Defocus range (μm) | 0.6 - 4.0 |
| Number of micrographs collected | 12 908 |
| **Cryo-EM reconstruction** |  |
| Particles in final reconstruction | 205 897 |
| Extracted particle box size (pixels) | 480 |
| Resolution (corrected, FSC 0.143) | 2.9 |
| Sharpening B-factor (Å<sup>2</sup>) | -109.4 |
| **Model composition** |  |
| Non-hydrogen atoms | 35 592 |
| Amino acid residues | 4 392 |
| Other ligands (ATP) | 6 |
| **Model statistics and validation** |  |
| Model deposition code | 9FP6 |
| Map-to-model FSC of 0.5 (Å) | 3.0 |
| Bond length RMS deviations (Å) | 0.002 |
| Angle RMS deviations (°) | 0.452 |
| MolProbity score | 1.51 |
| Clash score, all atoms | 6.19 |
| Ramachandran favoured % | 97.8 |
| Ramachandran outlier % | 0.0 |
| Rotamer outlier % | 1.4 |
| Cβ outliers % | 0.0 |

**Movie S1: Conformational changes undergone by NbNRC2 protomers transitioning from the resting state homodimer form to the activated homohexamer.**

**Data S1: List of modelled NLRs with their respective metadata and model statistics.**

**Data S2: List of contact residues from the cryo-EM model and AlphaFold 3 predicted structure.**

## Notes

https://doi.org/10.5281/zenodo.11546022

